# Intron-mediated induction of phenotypic heterogeneity

**DOI:** 10.1101/2021.01.19.427159

**Authors:** Martin Lukačišin, Adriana Espinosa-Cantú, Tobias Bollenbach

## Abstract

Introns are universally present in the nuclear genomes of eukaryotes^1^. The budding yeast, an otherwise intron-poor species, preserves two sets of ribosomal protein (RP) genes differing primarily in their introns^2–4^. Despite recent findings on the role of RP introns under stress and starvation^5–7^, understanding the contribution of introns to ribosome regulation remains challenging. Here, combining isogrowth profiling^8^ with single-cell protein measurements^9^, we found that introns can mediate inducible phenotypic heterogeneity conferring a clear fitness advantage. Osmotic stress leads to bimodal expression of the small ribosomal subunit protein Rps22B, mediated by an intron in the 5’ untranslated region of its transcript. The two resulting yeast subpopulations differ in their ability to cope with starvation. Low Rps22B protein levels resulted in prolonged survival under sustained starvation, while high Rps22B levels enabled cells to grow faster after transient starvation. Further, yeast growing at high sugar concentrations – similar to those in ripe grapes – exhibit bimodal Rps22B expression when approaching stationary phase. Differential intron-mediated regulation of RP genes thus provides a way to diversify the population when starvation looms in natural environments. Our findings reveal a new role for introns in inducing phenotypic heterogeneity in changing environments and suggest that duplicated RP genes in yeast contribute to resolving the evolutionary conflict between precise expression control and environmental responsiveness^10^.

## Main

Free-living cells are frequently challenged by changes in their environment, resulting in a need to allocate their resources accordingly. One of the most important cellular processes to manage is the production of ribosomes^11–13^. Ribosome synthesis, while essential for growth and division, consumes a substantial fraction of cellular resources^14,15^. To assemble into functional complexes, coordinated expression of tens of ribosomal RNA and ribosomal protein (RP) genes is necessary^16^. In the budding yeast, which underwent a whole genome duplication followed by loss or divergence of most duplicated genes^17,18^, RPs are largely conserved as duplicates (ohnologs) with high sequence identity^3^. Why evolution may have favoured the retention of RP ohnologs remains an open question; possible reasons include increased gene dosage, genetic robustness towards mutations, and distinct biological roles or differential regulation of the duplicated genes^3,17,19-27^. Recently, it has been suggested that duplicated transcription factors, which are also preferentially retained, might be an evolutionary solution to a trade-off between tight regulation of expression level and responsiveness to environmental changes, such as stress^10^. In this scenario, one ohnolog provides a precise level of protein expression required irrespective of the environment, while the other one generates population heterogeneity, allowing for a flexible response when the environment changes. While the phenotypic effects of deleting the less-expressed copy of duplicated ribosomal genes are usually observed only under stress^21^, it is unclear if this paradigm applies to duplicated ribosomal proteins.

The duplicated RP genes in yeast are highly enriched for introns. While fewer than 5% of all yeast genes contain an intron^28^, 94 out of 118 (80%) RP genes with an ohnolog do^4^. Differential expression of RPs through intronic regulation could expand their functional repertoire, even within the low sequence divergence; hence, introns may be involved in the evolutionary conservation of RP ohnologs^6,22^. Yeast introns in RP genes affect the expression level of the corresponding gene and, in some instances, that of its ohnolog^7,22,29-31^. Concerted intron retention in transcripts occurs in response to stress, suggesting a functional role for splicing regulation in yeast^32^. Such a role is supported by the observation that intron deletion in the budding yeast results in growth alterations under conditions including drug treatment, starvation, or population saturation^5–7^. Introns are thus clearly relevant for adapting to environmental changes; nevertheless, the determinants and functional outcomes of intron-mediated responses remain enigmatic.

Since ribosome synthesis and protein translation are tightly coupled to growth rate^33^, it is crucial to dissect cellular responses that are specific to particular stressors from responses to non-specific growth-inhibition when studying the effects of stress on ribosomal function. To this end, we have recently established a method termed isogrowth profiling^8^. It is based on exposing cells to a combination of two drugs in an antiparallel concentration gradient discretized into separate liquid cultures, while keeping the overall inhibition constant. Analyzing the transcriptome along the growth isobole then enables distinguishing responses specific to each drug, or to their combination, from the general stress and growth inhibition responses.

Here, to explore the role of introns and RPs in stress response, we extend isogrowth profiling from RNA sequencing to single cell protein-level measurements. We found that lithium chloride (LiCl) induces extensive intron retention in RP transcripts and an intron-dependent bimodal expression of Rps22B, a component of the small ribosomal subunit. The two subpopulations exhibit differential fitness under starvation and recovery. We show that while yeast in standard rich laboratory growth medium do not exhibit Rps22B bimodality, cells in medium with high glucose concentration do as they approach stationarity. Together, these results suggest that yeast has evolved an intron-mediated regulation mechanism of Rps22B to cope with uncertainty regarding possible replenishment of nutrients at the end of exponential growth in a high-glucose environment.

### LiCl inhibits splicing of ribosomal protein transcripts

To study the effect of stress on the regulation of ribosomes, we started by investigating the transcriptional response to two growth inhibitors – LiCl, a pleiotropic drug inducing cationic and osmotic stress^34^, and cycloheximide, an inhibitor of the large ribosomal subunit^33^. Since regulation of ribosomes is sensitive to changes in growth rate and medium composition^33^, we used an antiparallel concentration gradient of the two drugs applied in separate liquid cultures^8^, which consistently lowered the relative growth rate to near 50% (**Fig. 1A-B**). We inoculated the drug-free control at a lower cell density so that all samples reach comparable cell density at the time of harvesting (**Fig. 1B**). To determine intron retention, RNA was extracted from the samples, ribo-depleted and sequenced (*Methods*). This procedure on the series of drug-mixtures enabled us to observe the specific effects of stressors on intron retention in a way that is not confounded by growth-rate or cell-density effects.

**Figure 1:**
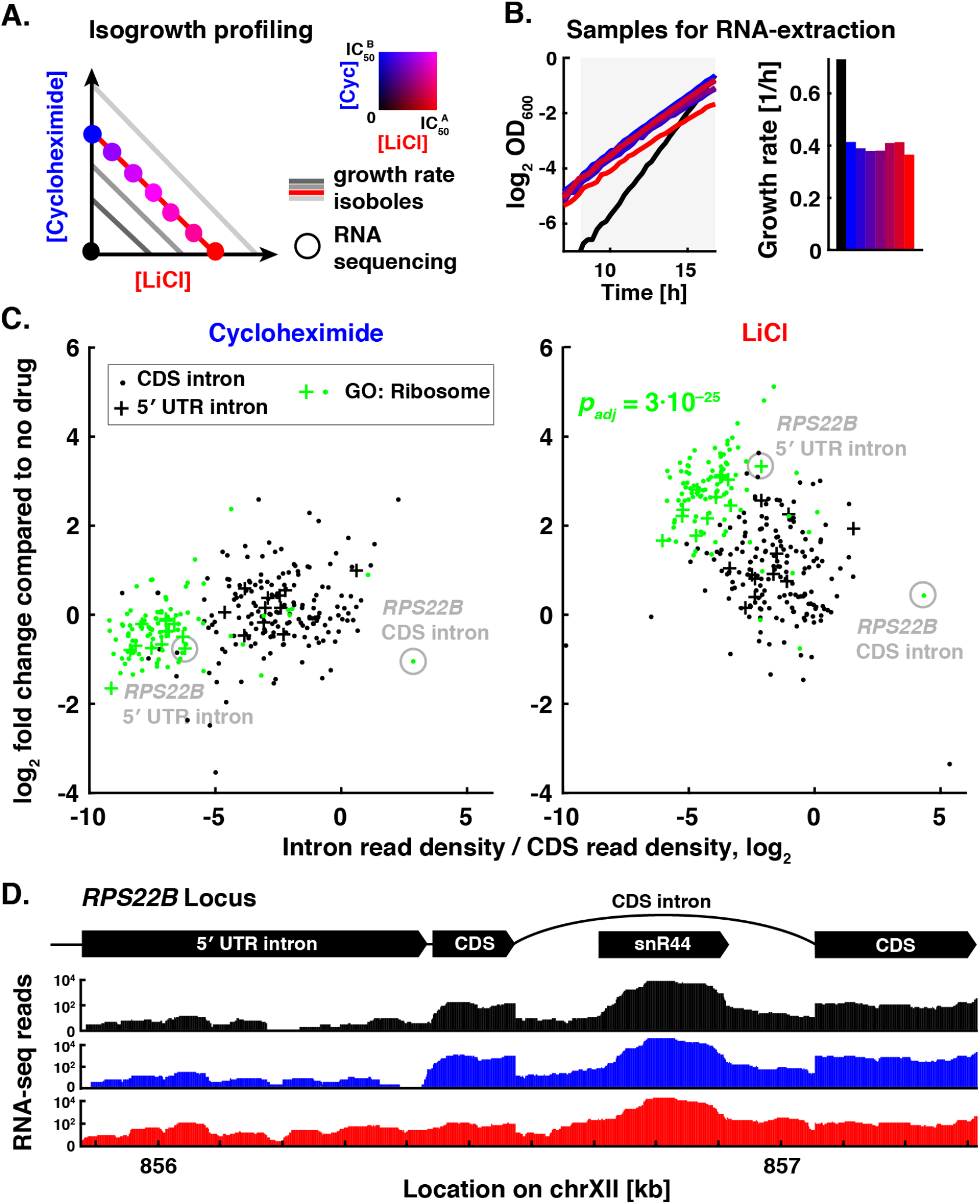
LiCl stress inhibits mRNA splicing of ribosomal proteins. **A.** Schematic of isogrowth profiling used to characterize changes of splicing in mRNA. The colored dots indicate the points in the two-drug gradient used to extract total RNA that was subsequently ribo-depleted and sequenced. **B.** Growth curves of samples used for RNA-sequencing. The no-drug control was inoculated at lower density to reach comparable optical density at the time of extraction. Shaded area denotes measurements used to determine the exponential growth rates (right). Color code as in (A). **C.** Intron retention rate (intron read density / CDS read density, *Methods*) for all nuclear introns when treated with cycloheximide or LiCl, normalized to no-drug control (y-axis) vs. non-normalized (x-axis). The most significantly enriched GO Cellular Component gene set for the LiCl sample (‘ribosome’) is displayed in green with the corresponding *p*-value of a hypergeometric test. CDS introns are represented by dots; 5’ UTR introns by plus signs. See **Fig. S1** for data at other points of the isobole. **D.** RNA sequencing counts for the *RPS22B* locus, color coded as in A. Above: Schematic of the *RPS22B* introns and exons and *snR44* locus according to Saccharomyces Genome Database.

We found that LiCl induces extensive intron retention compared to the drug-free control (**Fig. 1C**). Intron retention was not due to changes in growth rate or general stress, as no such increase was observed when cells were treated with cycloheximide (**Fig. S1A**), or two other drugs with different targets (**Fig. S2A**). RP transcripts were the functional gene set most strongly affected by this increase in intron retention (BH-corrected hypergeometric test, *p_adj_* = 3·10^-25^, **Fig. 1C** and **S1B-C**; *Methods*). The increase in intron retention was correlated with a decrease in their host transcript level (**Fig. S1D-E**). Analyzing RNA-sequencing reads that span the exon-intron junctions confirmed that introns are retained in mature transcripts, rather than being spliced out but not degraded (**Fig. S1F**); this observation was further corroborated by observing the increase in intron retention also in the polyadenylated fraction of RNA (**Fig. S2B**). It has been previously proposed that splicing of RP genes is downregulated by accumulation of stable excised introns in the stationary phase^5^. However, these introns were not retained differently to other introns under LiCl stress (**Fig. S1G**), suggesting that the high intron retention in RP transcripts observed under LiCl is not mediated by the stable linear excised introns previously reported.

### Osmotic stress induces bimodal expression of Rps22B

*RPS22B* was a clear outlier with respect to intron retention under LiCl treatment. *RPS22B* encodes a component of the small ribosomal subunit and is one of only nine *S. cerevisiae* genes containing two introns^36^. The intron in the 5’ untranslated region (5’ UTR) of *RPS22B* showed the highest increase in intron retention among 5’ UTR introns under osmotic stress (**Fig. 1C**), while there appeared to be numerous copies of the coding sequence (CDS) intron of *RPS22B*, presumably due to a small RNA encoded within the intron that is under separate regulation (**Fig. 1D**, *snR44*). The prevalence of stop codons in the introns of *RPS22B* (**Fig. S3A-B**), typical of yeast introns^39,40^, suggested that intron retention in the transcript could manifest itself in altered protein levels rather than protein isoforms. To observe the single-cell behaviour of Rps22B at the protein level, we coupled the rigorously controlled combinatorial drug treatment to flow cytometry using a yeast strain with a GFP-tagged Rps22B^9^. Notably, Rps22B exhibited two clearly distinct expression levels at intermediate LiCl concentrations (**Fig. 2A**). Other osmotic stressors similarly induced bimodal expression of Rps22B (**Fig. S3C**).

**Figure 2:**
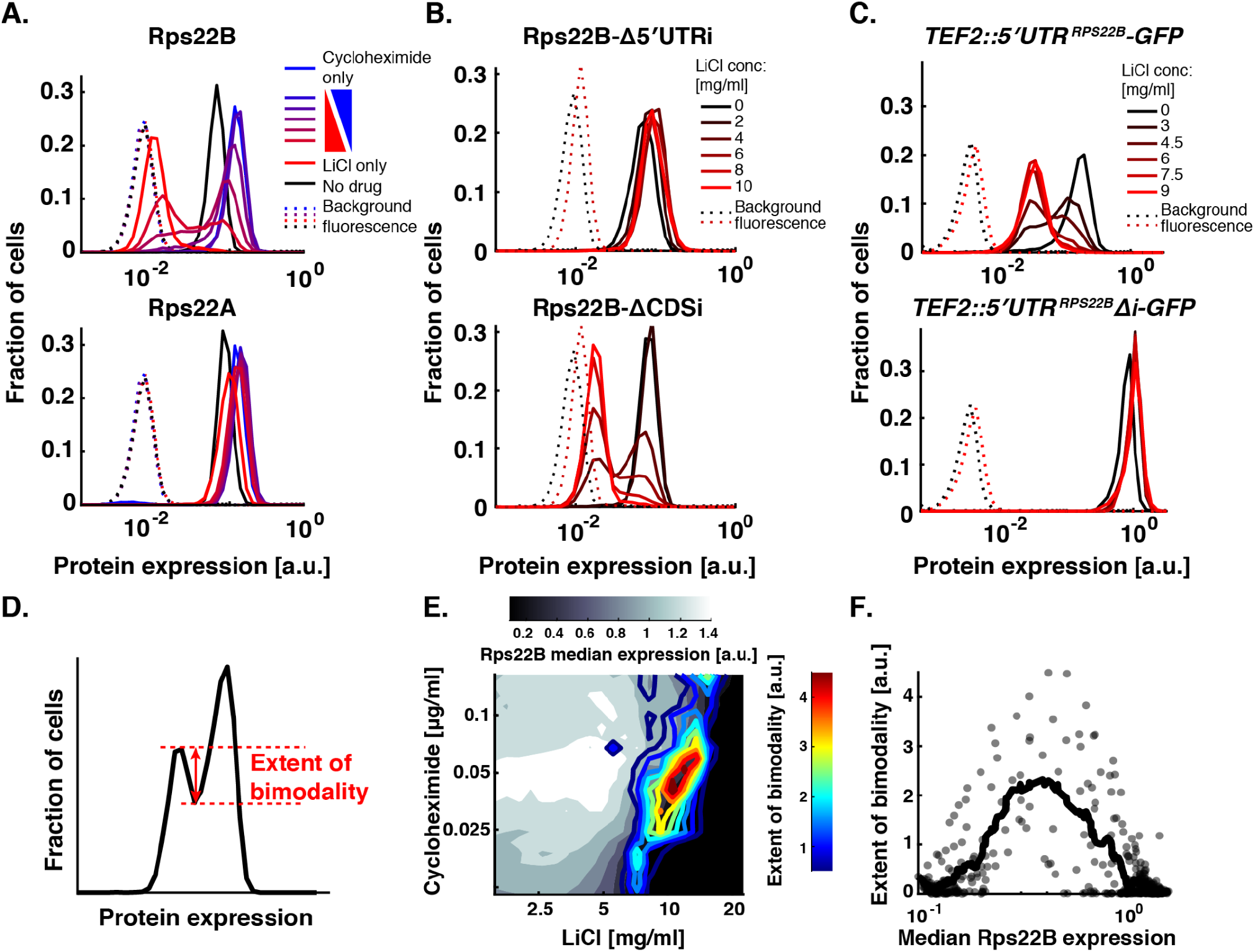
5’ UTR intron mediates bimodal Rps22B protein expression under LiCl, consistent with a bistable regulatory loop. **A.** Histograms of flow cytometry measurement of strains with GFP-tagged Rps22B or Rps22A. The ribosomal protein gene *RPS22B*, containing the 5’ UTR intron with largest increase in retention due to LiCl, exhibits bimodal protein expression at intermediate LiCl concentrations, whereas its ohnologue, *RPS22A* that contains no introns, does not. See **Fig. S3** for Rps22B expression in other osmotic stresses. **B.** As (A), but with seamless deletion of either of the *RPS22B* introns. See **Fig. S3D** for a strain with both introns deleted. **C.** As (A), but for strains with 5’ UTR of *RPS22B* fused to *GFP*, with or without the 5’ UTR intron. **D.** Ad hoc definition of the extent of Rps22B bimodality (*Methods*). **E.** The extent of Rps22B bimodality (coloured lines) is overlaid on Rps22B median protein level (greyscale) in a 2-drug gradient of LiCl and cycloheximide. **F.** Extent of Rps22B bimodality as a function of median Rps22B protein level is shown for all wells in the 2-drug gradient (dots) alongside the running average with a window of 30 data points (line). Rps22B bimodality peaks at a certain level of median Rps22B expression, rather than at a certain growth rate or LiCl concentration. See **Fig. S4 and S5** for comparison with LiCl-myriocin drug pair.

### 5’ UTR intron mediates Rps22B bimodality

While LiCl induced bimodal expression of Rps22B, no such expression pattern was observed for Rps22A, the intronless ohnologue of Rps22B (**Fig. 2A**), whose protein sequence is identical to that of Rps22B except for a single amino acid (**Fig. S3D**). Since the promoters of *RPS22A* and *RPS22B* behave similarly under different conditions^39^, we hypothesized that the bimodal expression of Rps22B observed under osmotic stress depends on the presence of one or both introns; especially the 5’ UTR intron is remarkably conserved both within *Saccharomycetaceae*^28^ and *S. cerevisiae* itself (**Fig. S3E**). Deletion of the 5’ UTR intron, but not the CDS intron, abrogated the bimodal expression of Rps22B under LiCl stress (**Fig. 2B and S2F**). In addition, it resulted in higher levels of *RPS22B* transcripts (**Fig. S3G**) and protein (**Fig. 2B and S3F**) under LiCl stress compared to the parental strain. No growth defects that could explain the differences in protein expression between the strains were observed (**Fig. S3H**). This suggests that, in the absence of the 5’ UTR intron, cells under LiCl stress can no longer downregulate Rps22B production, consistent with previous reports that the dsRNA structure contained in the intron is necessary for the regulated degradation of the transcript^40^. Fusing the 5’ UTR of *RPS22B* to *GFP* conferred bimodal GFP expression under LiCl stress, but only if the 5’ UTR intron was left intact (**Fig. 2C**). These results show that the 5’ UTR of *RPS22B* is a cis-regulatory element sufficient to confer bimodal protein expression to an unrelated gene and corroborate that the evolutionarily conserved 5’ UTR intron is necessary for this effect.

To further elucidate the role of the 5’ UTR intron in the protein heterogeneity of Rps22B, we quantified intron retention in different situations. Fluorescence-activated cell sorting (FACS) of the LiCl-induced Rps22B-bimodal cells showed that Rps22B-high cells have lower intron retention and higher CDS transcript level; however, these differences alone appear insufficient to account for the pronounced protein expression bimodality (**Fig. S3G**). Moreover, sequencing the polyadenylated RNA fraction revealed that while LiCl induces a strong increase in 5’ UTR intron retention in the Rps22B transcript, this increase is not necessary for the induction of Rps22B protein bimodality by other osmotic stressors (**Fig. S3G**). This suggests that the bimodal expression of Rps22B under osmotic stress is mediated by a post-transcriptional mechanism that requires the presence of the 5’ UTR intron and amplifies cell-to-cell differences in intron retention at the protein level.

### Rps22B expression pattern is consistent with a bistable regulatory loop

A bimodal expression pattern suggests – but does not necessarily imply – an underlying bistable regulatory circuit, such as a positive feedback loop^41,42^. A general hallmark of such a regulatory circuit is the existence of an unstable fixed point. Here, this fixed point would correspond to a protein level at which a small increase or decrease causes the cell to go to one of the two stable fixed points corresponding to high or low expression states^43^, respectively. To address if such an unstable fixed point may exist for Rps22B, we explored the Rps22B bimodality in the presence of LiCl stress while using another drug to perturb the growth rate and, as measured, the overall level of Rps22B. We used an automated liquid handling setup that enables continuous monitoring of culture over a fine two-drug gradient distributed over six 96-well microtitre plates^8^ (**Fig. S4A**) and at the end of the incubation measured the single-cell protein expression by flow cytometry (*Methods*). We used LiCl in combination with two drugs with disparate mechanisms, the translation inhibitor cycloheximide and the sphingolipid synthesis inhibitor myriocin^44^. We found that the extent of how clearly the expression levels of the two subpopulations are separated into two peaks depends primarily on the median protein level of Rps22B (**Fig. 2D-E and S4C-D**), with a maximum at an intermediate level (**Fig. 2F and S4E**). This behaviour is consistent with the existence of an unstable fixed point at this protein level, causing what would be an otherwise log-normally distributed population centered around this point to divide equally into the two presumably stable subpopulations.

### Single-cell isogrowth profiling uncovers environment-induced protein heterogeneity

To explore the extent to which osmotic or translation stress induce distinct cellular states, as observed for Rps22B, we took advantage of the yeast protein-GFP library^9^ containing 4,156 strains with single protein-GFP fusions and performed genome-wide single-cell isogrowth profiling (**Fig. S6A-C**). We profiled the entire library in four of the conditions used previously (**Table S1**) and selected strains which, on visual inspection, did not show a clearly unimodal expression pattern, for further characterization using a more detailed antiparallel gradient (**Table S2**). We found several instances of protein expression heterogeneity resulting from non-specific growth inhibition (**Fig. S6D**) or specifically from either of the stresses (**Fig. S6E**). Cycloheximide induced bimodal expression of Hsp12, the budding yeast persistence marker^45^, and of Aro9, an enzyme involved in production of yeast quorum sensing molecule tryptophol^46–48^, a trigger for invasive growth in low-nitrogen environment^47^. LiCl induced heterogeneity in Rps9A, another small ribosomal protein subunit containing an intron in its gene. Similar to Rps22B, the stress-induced heterogeneity of Rps9A levels was intron-mediated (**Fig. S6F**). Additionally, there was a general trend in which an increase in intron retention in LiCl correlated with a decrease in the level of the respective protein (**Fig. S6G**). Overall, these observations indicate that intron retention is used by the yeast cell to control the protein level and, in the case of Rps9A and Rps22B, also the protein level heterogeneity.

### Rps22B bimodality leads to differential fitness in starvation

Introns play a key role in preparing the yeast population for starvation^6,32^. Therefore, we hypothesized that the two subpopulations, defined by distinct levels of Rps22B expression, possess differential fitness under starvation stress. To test this idea, we subjected an exponentially growing culture of the Rps22B-GFP strain to LiCl stress using a microfluidic system (*Methods*). Following the establishment of bimodality, we replaced the LiCl-containing growth medium with spent medium. In this way, we induced starvation of varying duration, followed by a sudden switch to rich medium (**Fig. 3A**). At first, the spent medium triggered a rapid disappearance of the strong nucleolar signal of Rps22B-GFP (**Supplementary Movies M1 and M2**), presumably caused by the redistribution of nucleolar proteins across the cytoplasm upon stress, as occurs in other model systems^49^. Following the switch to rich medium, the nucleolar signal of Rps22B re-emerged in cells that started growing again. Notably, the fitness of cells depended on their Rps22B expression level immediately preceding the starvation stress. Longer starvation stress favoured the survival of Rps22B-lowly expressing cells in that they lysed less frequently than the highly-expressing cells. In contrast, shorter starvation favoured the Rps22B-highly expressing cells in that they budded more in the period following nutrient replenishment (**Fig. 3B-F**). FACS corroborated this phenotypic difference, as the Rps22B-GFP-high subpopulation exhibited poorer survival under starvation (**Fig. 3H**), but higher growth rate in rich medium (**Fig. 3I-J**) compared to the Rps22B-GFP-low subpopulation. This marked diversification of the population observed for Rps22B, with clear fitness effects under a subsequent stress may reflect a bet-hedging strategy^50–52^.

**Figure 3:**
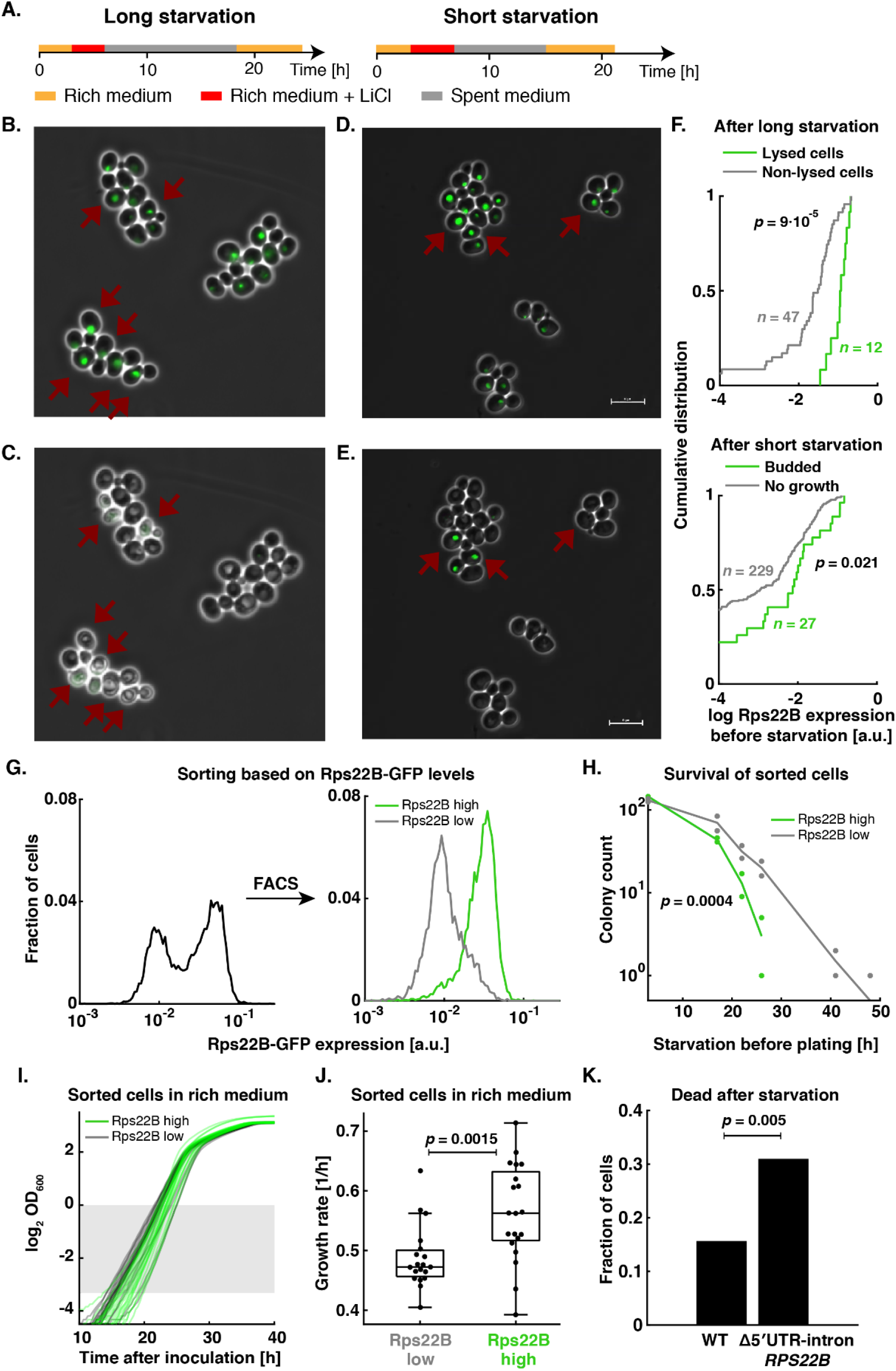
Rps22B expression level confers selective advantage on cells under starvation stress of varying duration. **A.** Schematic of the temporal sequence of media used in microfluidic microscopy experiments with the Rps22B-GFP strain. **B-C.** Example micrograph of Rps22B-GFP yeast cells after application of LiCl, showing LiCl-induced bimodality of Rps22B expression (B). The Rps22B-high cells (dark red arrows) lyse following prolonged starvation and nutrient replenishment (C). **D-E.** As (B-C), but for shorter starvation stress. Following a short starvation stress and medium replenishment, the Rps22B high expressing cells marked with dark red arrows (D) budded more than other cells (E). Scale bar = 10 μm. See time-lapse **Supplementary Movies 1 and 2**. **F.** Quantification of time-lapse micrographs from microfluidic starvation experiments. Upper panel: Cumulative distribution of Rps22B expression levels measured before starvation in cells that have either lysed (green) or not lysed (grey) within 6 h after medium replenishment following a long starvation. Lower panel: Cumulative distribution of Rps22B expression levels before starvation in cells that have either budded (green) or not budded (grey) within 6 h after medium replenishment following a shorter starvation. Rps22B expression values between the two panels are not comparable. Mann-Whitney *U* test was used to determine significance. **G.** Histograms of Rps22B-GFP expression in LiCl before and after fluorescence-activated cell sorting. **H.** Survival curves (colony-forming units) of the sorted populations as a function of time spent in PBS following the sorting and prior to plating on rich medium. Significance was determined by bootstrapping (*Methods*). **I.** Growth curves of cultures in rich medium following sorting. Shaded rectangle denotes OD range used to quantify growth rates. **J.** Quantification of growth rates from (I). Significance was determined using two-sided *t*-test. **K.** Fraction of cells dead after starvation in time-lapse microscopy experiments comparing WT and *RPS22B* 5’ UTR intron deletion mutant. For experimental setup and further quantification see **Fig. S7**. Significance was determined using a permutation test.

To test whether the link between intron-mediated Rps22B protein heterogeneity and phenotypic heterogeneity is causal, we performed time-lapse microscopy with the *RPS22B* 5’ UTR intron deletion mutant (**Fig. S7A**). Deletion of the 5’ UTR intron in *RPS22B* not only increased the fraction of Rps22B-GFP high cells, but also the fraction of cells dying under starvation (**Fig. 3K, S7B-D**) and largely abolished the phenotypic heterogeneity associated with Rps22B protein heterogeneity (**Fig. S7E**). Population-level assays with the intron-deletion mutant confirmed that its survival times under starvation are less heterogeneous and lower on average (**Fig. S8A**) while its growth rate is increased under LiCl stress (**Fig. S8B**). These observations confirm that not only the Rps22B expression heterogeneity, but also the phenotypic heterogeneity is mediated by the *RPS22B* 5’ UTR intron.

### Yeast nearing saturation in high-glucose medium exhibit bimodal Rps22B expression

Bet-hedging strategies can evolve if the eliciting signal is probabilistically followed by stress in which the subpopulations exhibit differential fitness in the natural environment^50,53^. However, yeasts nearing saturation, and hence starvation, in standard rich laboratory medium do not exhibit Rps22B bimodality (**Fig. 4A**). Therefore, we wondered if there is a plausible natural setting in which Rps22B bimodality is triggered just before the onset of starvation. We reasoned that yeasts may often be exposed to hyperosmotic sugar concentrations, such as those in ripe fruits. High glucose elicits osmotic responses that are different from those to salt^54^. Additionally, high glucose concentration, despite being an osmotic stressor, leads to comparatively higher expression levels of ribosomal proteins than LiCl stress (**Fig. S9**). Since Rps22B exhibits the maximum extent of bimodality at a certain median expression level (**Fig. 2D-F**), we hypothesized that a high-glucose environment might specifically induce Rps22B bimodality as yeast nears saturation, when ribosomal protein levels decrease due to growth slowdown.

**Figure 4:**
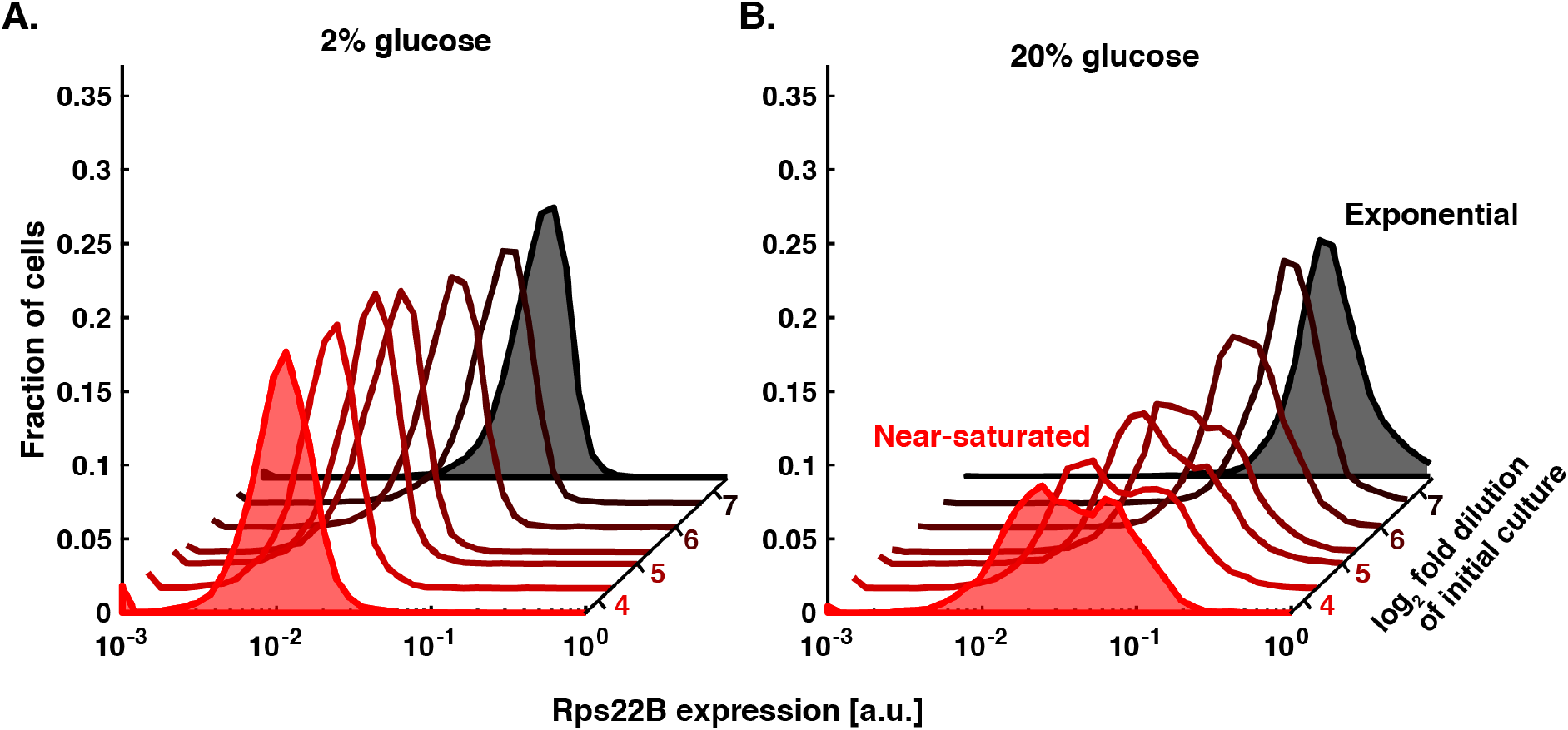
High-glucose environment induces bimodal Rps22B expression as cells near saturation. Histograms of Rps22-GFP expression as measured by flow cytometry after 10 h of incubation in standard rich medium (**A**), or in yeast extract-peptone medium supplemented with 20% (w/v) glucose (**B**). Red hue indicates increasing cell density in the initial inoculum. Examples of exponential and near-saturated cultures are highlighted by shading.

To test this idea, we inoculated Rps22B-GFP cultures into a yeast-peptone medium with high glucose concentration (20% w/v) that emulates the total hexose concentration in ripe grapes^55^, at various initial cell densities. Exponentially growing cells in high glucose medium generally showed less clear bimodal expression of Rps22B-GFP than in LiCl; however, cultures in high glucose medium that were nearing saturation exhibited clearly bimodal Rps22B expression (**Fig. 4B**). This observation suggests that part of the yeast population is attuned to a probabilistic event of nutrient replenishment that can follow growth in a high-glucose environment, while the rest of the population is preparing for starvation.

## Discussion

In the budding yeast, the overall expression level of ribosomal proteins results from two sets of ribosomal protein genes with high amino-acid sequence similarity, often differing in the presence and identity of intronic sequences^7,22^. Here, we uncovered an intron-mediated regulation of protein expression heterogeneity. We showed that LiCl leads to a widespread retention of introns^56^ that is independent of growth rate perturbations and predominantly affects ribosomal protein transcripts. The small ribosomal subunit protein gene *RPS22B* manifested bimodal protein expression in osmotic stress, mediated by its 5’ UTR intron, which is conserved throughout the *Saccharomycetaceae*^28^. In contrast, its intronless ohnolog *RPS22A*, exhibited a unimodal protein expression level irrespective of stress. This behaviour of the *RPS22* gene pair offers a new paradigm to explain the function of introns in duplicated ribosomal protein genes, in that they enable differential and versatile regulation for duplicated genes as an evolutionary trade-off between environmental responsiveness and precise regulation^10^.

We found that population diversification due to bimodal expression of Rps22B leads to differential fitness of these subpopulations in the face of starvation stress and subsequent recovery. Reminiscent of bet-hedging strategies^50^, we found that in high glucose medium, yeast populations diversify in Rps22B expression as they enter stationary phase, as if anticipating a probabilistic replenishment of nutrients. Why should such a behaviour have evolved for a high-glucose environment, but not manifest at glucose levels present in the standard laboratory “rich” medium? One plausible scenario is that for yeast living on the skin of grapes, high glucose concentration is an environmental signal that the fruits are ripening and bursting. Osmotic bursting of fruits is known to follow a probabilistic trajectory over time^57^ and is known to happen predominantly during rainfall^58^; hence, in a cluster of grapes, vigorous yeast growth due to bursting of one of the berries might be followed by nutrient replenishment due to bursting of a neighbouring berry, determined by the probabilistic aspects of rain duration and intensity.

The fact that we found osmotic stress-induced, intron-mediated phenotypic heterogeneity only for Rps22B and Rps9A, while most other pairs of duplicated ribosomal protein genes also contain at least one intron, is intriguing, especially since the effect of introns on protein expression level has been reported for other ribosomal proteins in the budding yeast^7,59^. It is thus tempting to speculate that for other ribosomal proteins, there might exist other levels of osmotic stress, or other stressors altogether, that would trigger population diversification with respect to the expression level of a given ribosomal protein. Such a set of diversification mechanisms could then present a versatile stress toolkit for the yeast population bracing against continued stress, at the same time maintaining a small, stress-sensitive subpopulation poised to rapidly exploit a short window of fitness advantage should the stress suddenly disappear. Our study thus highlights the need of studying the intronic regulation of population diversification and demonstrates the utility of using graded, growth-rate controlled perturbations.

## Methods

### Transcriptional isogrowth profiling

For isogrowth RNA sequencing, *S. cerevisiae* strain BY4741 was grown in 7 conditions of varying ratios of LiCl (Sigma Aldrich, L9650) and cycloheximide (Sigma Aldrich, 37094), ensuring 50% growth inhibition, and in YPD [yeast extract (Sigma Aldrich cat. No. Y1625) 1% w/v, peptone (Sigma Aldrich cat. No. 91249) 2% w/v, dextrose (Sigma Aldrich cat. No. D9434) 2% w/v] containing no drug (Table 1). The frozen stock was diluted 130-times into drug-containing wells and 100-times more into wells without drug. The cells were incubated for a total of ~17 h to a final absorbance of ~0.1 on the 96-well plate, corresponding to OD_600_~0.5. Cells were then harvested and the RNA extraction was performed using RiboPure RNA Purification Kit for yeast (Thermo Scientific, AM1926). The extracted RNA was sent to the Next Generation Sequencing Facility of the Vienna Biocenter Core Facilities, where it was rRNA depleted using Illumina Ribozero Yeast Kit, multiplexed using Illumina True Seq adapters, single-end 50bp sequenced using Illumina HiSeqV4 and demultiplexed.

**Table 1:** Concentrations of drugs used for the RNA-isogrowth profiling. LiCl concentration is stated both in weight and molar concentration for an easier comparison to published literature.

| Condition<br>(as labeled in Fig. S1) | Cycloheximide<br>[μg/ml] | LiCl<br>[mg/ml] | LiCl<br>[mM] |
| --- | --- | --- | --- |
| 1 | 0.11 | 0 | 0 |
| 2 | 0.095 | 1.2 | 28 |
| 3 | 0.083 | 2.5 | 59 |
| 4 | 0.069 | 4.1 | 97 |
| 5 | 0.049 | 5.9 | 139 |
| 6 | 0.026 | 7.7 | 182 |
| 7 | 0 | 8.5 | 201 |

RNA samples from treatments with single stressors were obtained from cultures with automated re-inoculation every 8 hours^8^, with a total incubation time of 24 hours, to keep cultures in exponential phase and assure the drug effects have taken place. Strains were grown in YPD with a drug, either in a concentration gradient to select samples closest to 50% of growth inhibition (LiCl, Fenpropimorph, and Myriocin), or in a single concentration (0.4 M NaCl and glucose 20%). Fenpropimorph (Sigma Aldrich cat. No. 36772) was tested in a 0.5-1.5 μM gradient, sequenced samples were treated with 0.85 μM Fenpropimorph. Myriocin (Sigma Aldrich cat. No. 476300) gradient encompassed 0.25-0.75 μg/ml; 0.32 μg/ml Myroicin samples were sequenced. For each sample of extracted RNA, both ribodepleted total RNA and mRNA Seq libraries were prepared at the Cologne Center for Genomics (CCG) and sequenced using 2×100bp paired-end reads.

### Intron retention analysis

The reads resulting from sequencing were aligned to the annotated reference *S. cerevisiae* genome R64-2 using TopHat^60^ or HISAT2^61^. The reads mapping to introns or CDS of intron-containing genes were quantified using featureCounts^62^ using custom produced .saf files extracted from the reference .gff annotation file. The intron retention rate was calculated as the ratio of intron counts normalized to the intron length over the CDS counts normalized to the length of the CDS region. For genes containing multiple introns, the introns were treated jointly for CDS introns and separately for 5’ UTR introns. Alignments were visualized using Integrative Genomics Viewer^63^.

For the analysis shown in **Fig. S1F**, first the number of contiguous reads (no mismatches allowed) overlapping the individual intron ends was determined. This count was then averaged between 5’ and 3’-end of the intron ends and divided by the read length (50bp). The intron retention rate was then calculated as the ratio of the resulting value over the CDS counts normalized to the length of the CDS region.

For GO enrichment analysis, intron-containing genes were ordered into a ranked list based on the fold-change increase in intron retention in LiCl compared to the no-drug control, either in decreasing (**Fig. 1C**) or increasing order (**Fig. S1B**). The ranked list was then analysed using GOrilla^64^, for either Cellular Component or Biological Process, respectively. The adjusted *p*-values reported here correspond to the false discovery rate reported by GOrilla. Gene enrichments were visualized using Revigo^65^.

### Rps22B intron conservation analysis

To gauge the evolutionary conservation of the Rps22B 5’ UTR intron within the natural population of *S. cerevisiae* (**Fig. S3E**), previously published sequencing data for 1011 isolates^66^ were multiple-sequence aligned using Clustal Omega^67^ and visualised using MSA-BIOJS^68^.

### Screening the entire yeast protein-GFP library for drug-induced bimodality

The whole yeast protein-GFP library^9^, contained in 43 96-well microplates, was screened for drug-induced bimodality by culturing in four conditions: YPD medium containing no drug, cycloheximide, LiCl, or a combination of LiCl and cycloheximide. The screen was divided into 6 experimental batches. In every batch, up to 9 plates of the library were processed, with the same strain being cultured in four conditions on the same day and on the same microtitre plate, giving rise to up to 36 microtitre plates per experiment. Before the experiment, empty microtitre plates and YPD medium were pre-incubated at 30°C to ensure reproducibility of gene expression measurements. LiCl, cycloheximide, or a mixture of the two, were diluted with YPD to a final concentration needed to achieve 40% inhibition and ensuring that in the mixed condition, the LiCl and cycloheximide were mixed in an equipotent manner, meaning that on their own they both elicited approximately the same (smaller) growth inhibition (Table 2). The drug solutions were pipetted row-wise into a 96-well microtitre plate, including a drug-free YPD control, enabling the cultivation of 24 strains in 4 conditions per microtitre plate. The library plates stored at −80°C were thawed and, using an electronic multi-step multi-channel pipette, 1.5 μl saturated glycerol stock of the corresponding strain was inoculated into each drug-containing well and 0.2 μl of the saturated glycerol stock was inoculated into drug-free YPD as a control. The smaller inoculum size for the drug-free control was designed to ensure that the final cell density at the end of the experiment is comparable to that of the drug-containing wells. Plates were incubated for ~14 h in an automated incubator (Liconic StoreX) kept at 30°C, >95% humidity, vigorously shaken at >1000 rpm. During the incubation, OD_600_ was measured every ~45 min in a Tecan Infinite F500 plate reader. In addition to shaking during incubation, directly before each measurement, plates were shaken on a magnetic shaker (Teleshake; Thermo Scientific) at 1100 rpm for 20 s. The growth rates were inferred by fitting a line to the log-linear part of OD_600_ measurements between 0.01 and 1.

**Table 2.** Concentrations of drugs used in the whole yeast protein-GFP library isogrowth screen. LiCl concentration is stated both in weight and molar concentration for an easier comparison to published literature.

| Condition | Cycloheximide [µg/ml] | LiCl [mg/ml] | LiCl [mM] |
| --- | --- | --- | --- |
| Cycloheximide only | 0.10 | 0 | 0 |
| Cycloheximide + LiCl | 0.050 | 6.3 | 149 |
| LiCl only | 0 | 13.5 | 318 |
| YPD only | 0 | 0 | 0 |

After incubation, the yeast cells in 96-well microtitre plates were twice: centrifuged at 1050 g for 3.5 min and resuspended in ice-cold Tris-EDTA buffer by vigorous shaking at 1000 rpm on a Titramax shaker for 30 s. After another centrifugation at 1050 g for 3.5 min, the cells were resuspended in 80 μl of Tris-EDTA and immediately stored at −80°C. On the day of the flow cytometry measurement, the plates were thawed on ice for ~3 h and kept on ice until the measurement. The fluorescence was measured for 10,000 cells using BD FACS Canto II equipped with a high-throughput sampler. The green fluorescence measured in FITC-H channel was normalized to the forward scatter FSC-H channel.

Strains exhibiting bimodality, minor peaks, or prominent changes in gene expression at visual inspection of the expression histograms were manually selected for a more detailed screen. The selected strains were re-streaked from the yeast protein-GFP library microtitre plates onto YPD-agar plates and single clones for each strain were picked and cultured in a 96-well microtitre plate to saturation. Glycerol was added to a final concentration of 15% and the plates were frozen. The plates were screened in 8 conditions and analysed using flow cytometry in a way analogous to that described above. The identity of the relevant strains – Rps22A, Rps22B, Aro9, Hsp12, Cit1 and Tdh1 – was confirmed by PCR and gel electrophoresis as reported before^9^, using a common F2CHK reverse primer and strain specific oligos taken from (https://yeastgfp.yeastgenome.org/yeastGFPOligoSequence.txt). The strain Smi1 appeared bimodal in the screen; however, its identity could not be confirmed as above. The identity of other strains was not tested.

### Construction of GFP-labeled intron deletion strains and flow cytometry experiments

The seamless intron deletion strains for Rps22B (Rps22B-Δi1-Δi2, rps22bDi_JPY138F4; Rps22B-Δi1, rps22bD1i_MDY125A4; Rps22B-Δi2, rps22bD2i_MDY125A9) and Rps9A (Rps9A-Δi, rps9aDi_MDY133H8), as well as the parental haploid strain WT_JPY10H3 (Mat**a** ura3Δ0 lys2Δ0 leu2Δ0 his3Δ200) were a kind gift of Julie Parenteau and Sherif Abou Elela^6^. The parental and intron deletion strains were labeled with GFP fused to the protein of interest following homologous recombination of the PCR-amplified fluorescent marker from the matching yeast protein-GFP library strains. To achieve this, the DNA of the Rps22B-GFP and Rps9A-GFP strains from the library was extracted using Yeast DNA Extraction Kit (Thermo Scientific, 78870). PCR Amplification of the GFP label with 40 base pairs flanking sequences was done using the corresponding F2 and R1 pair of primers from Huh et al (https://yeastgfp.yeastgenome.org/yeastGFPOligoSequence.txt) in the following reaction conditions: 2 ng/ml of DNA in a final mix of 20 μl, with 0.4 units of Phusion High-Fidelity DNA polymerase (Thermo Fisher Scientific, Cat# F530SPM), 4 μl of 5x Phusion HF buffer, 0.2 μM dNTPs, and 0.5 μM primers. PCRs were incubated for 3 min at 98°C, followed by 34 cycles of 98°C for 30 sec and 72°C for 2.5 min, and a final incubation at 72°C for 5 min. The resulting PCR products were transformed into the parental WT_JPY10H3 and Δi strains as in Howson et al.^69^, selecting on SD -HIS agar plates (Takara Bio, 630411; Diagonal GmbH&CoKG, Y1751). The correct genomic context insertion of GFP of single colonies was confirmed by PCR using a combination of primers which also allowed the confirmation of the intron deletion strain. To confirm the Rps22B-GFP fusions in Rps22B-Δi and the WT_JPY10H3 strains, we used primers rps22b-899F 5’-CCGTTATTCTTCTCGCAACC-3’ binding upstream the 5’ UTR intron and RPS22B-CHK^69^ 5’-ACTAGATGGTGTGATCGGGC-3’ binding in the CDS-intron sequence in combination with the reverse primer F2CHK^69^ 5’-AACCCGGGGATCCGTCGACC-3’, complementary to the GFP sequence. To confirm the Rps9A-GFP fusions in the Rps9A-Δi and WT_JPY10H3 strains, we used rps9a-800F 5’-GTTCGATTTCTTGGTCGGACGC-3’ upstream the open reading frame and F2CHK. Once the successful construction of the GFP reporter strains was confirmed, 20 μl of saturated overnight cultures were inoculated into 96-well microtiter plates containing 180 μl of YPD with the respective concentration of LiCl. The microtiter plates were incubated at 30°C and continuous shaking at 900 rpm on a Titramax shaker for 7 h, harvested and measured on a flow cytometer as described above. Strains with genome-integrated GFP with different 5’ UTR fusions were a kind gift from Gary Stormo’s lab^31^.

### Measurement of Rps22B expression in detailed 2D drug gradients

The re-inoculation setup, as reported previously^8^, in conjunction with flow cytometry measurements was used to measure Rps22B expression in detailed 2D gradients of LiCl and cycloheximide, and LiCl and myriocin (Sigma Aldrich, M1177). In brief, *Saccharomyces cerevisiae* strain with RPS22B gene fused to GFP protein from ORF-GFP library^9^, was grown in YPD broth in a conical flask overnight and then distributed into a 96-well plate. A customized robotic setup (Tecan Freedom Evo 150) with 8 liquid handling channels and a robotic manipulator was used to produce a two-dimensional discretised two-drug 24×24-well gradient in YPD spread over six 96-well plates and to inoculate the yeast overnight culture to final liquid volume in the well 200 μl and final absorbance 0.03 on the 96-well plate, corresponding to standardised optical density at 600 nm of 0.15. Working drug solutions were prepared either by adding the respective amounts of concentrated DMSO drug stocks thawed from −20°C storage (no refreezing) previously prepared from stock chemicals (cycloheximide and myriocin), or by dissolving directly in YPD and sterile-filtering (LiCl). The six plates were incubated for three iterations, each lasting ~8 h. After the incubation the cells were harvested and measured using flow cytometry as described above. Rps22B protein bimodality was quantified as the depth of the trough on the Rps22B protein single-cell expression histogram, that is the y-axis distance between the trough and the lower of the two peaks (**Fig. 2D**). To this end, Matlab function *findpeaks* was used to determine the prominence of the peak that was created from the trough through inversion of the histogram values.

### FACS of Rps22B-GFP and experiments with GFP-sorted cells

Constitutive cytosolic-mCherry expression was added to the parental and intron deletion GFP-fusion strains. To do so, a *TDH3::mCherry* construct was inserted in the HO locus by homologous recombination following transformation of a NotI-digested SLVA06 plasmid^70^, kindly provided by Michael Springer’s lab. The Rps22B-GFP strain with constitutive mCherry expression was inoculated into YPD and incubated at 30°C for 16 hours. The overnight culture was diluted 20-fold into 4.5 mg/ml LiCl in YPD, incubated for 6 h and washed twice with PBS. Positive mCherry cells were sorted into low-GFP and high-GFP populations with a Becton Dickinson INFLUX cell sorter. Around 8×10^6^ sorted cells of each condition were used for RNA extraction and sequencing (see Transcriptional isogrowth profiling, *Methods*). Samples of sorted cells at a final concentration of 4000 cells/ml were prepared with PBS for the following experiments. Immediately after sorting, 15 μl of the samples were inoculated into fresh YPD media in 24 replicates each. Optical density was measured every 20 min for 48 hours in a plate reader incubated at 30°C with constant shaking. OD was corrected by subtracting the background, and growth rates were inferred by a linear fit to the log-linear part of the corrected OD measurements between 0.02 and 0.2. Growth rates higher than 0.85 h^-1^,which resulted from bacterial contaminations, were discarded. A two-tailed *t*-test was performed to compare the growth-rates from the two sorted populations by using Matlab (ttest2). For scoring survival after starvation, sorted cells were incubated in PBS at 30°C and shaking at 200 rpm. At each sampling time point, 50 μl of starving cultures were spread onto an OmniTray plate (Thermo Scientific) filled with 35 ml of YPD-agar, in two replicates for each sorted population. Agar plates were cultivated at 30°C, pictures were taken after ~36 hours, and colonies were counted using ImageJ. To quantify the difference between low-GFP and high-GFP populations (**Fig. 3H**), we employed the difference Δ between the total low-GFP and high-GFP colony counts along all timepoints. To test the statistical significance of Δ, we used bootstrapping. In brief, we created 10^4^ surrogates (artificial datasets) complying with the null hypothesis that there is no difference between the populations. To create one such surrogate, we sampled – for each replicate and time point – the colony count from the Poisson distribution with a rate λ corresponding to the mean colony count for that time point. We determined the *p*-value as the fraction of surrogates which exhibit a higher Δ than the original experimental dataset.

### Time-lapse imaging of Rps22B-GFP strain during starvation

For micrographs shown in **Fig. 3**, the ORF-GFP library Rps22B-GFP strain was used; for experiments with the intron deletion mutants (**Fig. S7**), we used the 5’ UTR intron-deletion strain and its parental Rps22B-GFP strain, both transformed with a plasmid constitutively expressing mCherry (see FACS of Rps22B-GFP and experiments with GFP-sorted cells, *Methods*) to allow for easy tracking of cellular integrity, since cells lose the mCherry signal after lysis. Strains were inoculated into YPD at 30°C with shaking overnight. The resulting culture was diluted 20-fold and loaded to the CellASIC ONIX2 haploid yeast microfluidic plates (MerckMillipore) at 55 kPa pressure. A sequence of YPD, LiCl-YPD, starvation medium (spent medium or PBS), and YPD were flushed through the microfluidic chamber in a time-controlled manner at 10 kPa pressure. Spent medium was produced by incubation of the Rps22B-GFP strain in YPD medium for 7 days with subsequent sterile filtration. Starvation times were empirically selected such that an intermediate number of cells died in the long starvation condition or budded after the short starvation. The specimen was imaged every 5 min at multiple locations with Nikon Eclipse Ti inverted microscope using a 100× oil objective in a 30°C cage incubator. The micrographs of yeast from just before the starvation stress (**Fig. 3**) were segmented and fluorescence-quantified using CellProfiler^71^ and the fluorescence readout was log_10_-transformed. The time-lapse images were then visually scored to determine which cells budded/lysed after the replenishment of nutrients. The time-lapse movies from the experiments with the intron-deletion mutant were segmented and tracked using YeaZ^72^.

### *RPS22B* 5’ UTR intron-deletion mutant fitness experiments

The 5’ UTR intron-deletion strain and its parental strain, with Rps22B-GFP fusion and constitutive mCherry expression, were grown overnight in YPD. For growth rate determination experiments in liquid media, 5 ml of each saturated overnight culture was standardized to OD_600_ = 0.1 and inoculated in a 1/10 dilution into YPD containing 0, 2.25, 4.5, or 9 mg/ml LiCl in 12 replicates of each strain and condition. Optical density was measured every 5 min for 24 h in a plate reader during incubation at 30°C with constant shaking. OD and growth rates were processed as above (see FACS of Rps22B-GFP and experiments with GFP-sorted cells, *Methods*). For scoring survival under starvation, 1 ml of each overnight culture was inoculated into 19 ml of YPD with 9 mg/ml LiCl. Cultures were incubated at 30°C with 200 rpm shaking for 6 h. Cultures were standardized to OD_600_ = 1 and washed twice by centrifuging at 1050 g for 5 min and resuspending pellets in 20 ml of PBS. Dilutions of 5000 cells/ml were incubated at 30°C with constant shaking. Plating onto YPD-agar, incubation, and analyzing colony counts was done as explained above (see FACS of Rps22B-GFP and experiments with GFP-sorted cells, *Methods*).

### Rps22 bimodality in other osmotic stresses and in high glucose

The ORF-GFP library Rps22B-GFP cultures were inoculated into YPD medium, YPD containing 0.6M NaCl, 2M KCl (Sigma Aldrich, S3014 and P9541 respectively) or into a yeast-peptone medium with 5%, 10% or 20% (w/v) glucose (Sigma Aldrich, G8270). In order to measure fluorescence intensity of cells from different cell density and growth stages, 6 microtiter plates were prepared, each with the same conditions but with a gradient of initial inoculum sizes; in this way, each plate accounted for the stress array with different cell density and could be incubated and analyzed at different time points. To achieve this, saturated overnight cultures were diluted with YPD in a 4/5 serial dilution (i.e. 4ml were transferred to the final volume of 5ml) to obtain (4/5)^0^ to (4/5)^23^ dilutions. Subsequently, 15 μl of cell dilutions were inoculated in 185 μl of media. Plates were incubated in the Liconic StoreX and kept at 30°C, >95% humidity, with constant shaking. Plates were processed at 8, 12, 16, 20 and 24 hours after inoculation with an automated robotic system consisting of a ACell System (HighRes Biosolutions) integrated with a Lynx liquid handling system (Dynamic Devices), a plate reader (Synergy H1, BioTek), and a CytoFLEX flow cytometer (Beckman Coulter). First, plates were shaken and measured as described above to obtain the OD_600_ values. Cultures were then transferred to a new plate and, when necessary, diluted to an absorbance of ~0.3 with Tris-EDTA buffer in order to avoid high cell densities, and measured by flow cytometry.

## Data availability

The RNA sequencing data are available in Gene Expression Omnibus under accession no. GSE155060. The processed flow cytometry data are available as Supplementary Table 1 and 2 accompanying this manuscript.

## Acknowledgements

We would like to thank the IST Austria Life Science Facility, the Miba Machine Shop, and Marta Lukačišinová for the support of the liquid handling robot. The Bioimaging Facility at IST Austria, Jeffrey Power and Berenike Meier at the University of Cologne, and Christoph Göttlinger at the FACS Analysis Facility at the Institute for Genetics, University of Cologne, for the support of flow cytometry experiments. Lucas Horst for the development of the automated experimental methods in Cologne. Julie Parenteau and Sherif Abou Elela, Gary Stormo, Mike Springer, and Maya Schuldiner for providing us with critical yeast strains. Booshini Fernando, Theresa Fink, Gerrit Ansmann and Guillaume Chevreau for technical support. Hania Köver, Gašper Tkačik, Nick Barton, Andreas Angermayr and Bor Kavčič for support during lab relocation. Daria Siekhaus, Mike Springer, and all the members of the Bollenbach group for support and fruitful discussions. Karin Mitosch, Marta Lukačišinová, Gianni Liti, and Alexander de Luna for the critical reading of our manuscript. This work was supported in part by Austrian Science Fund (FWF) standalone grant P 27201-B22 (to T.B.), HFSP program Grant RGP0042/2013 (to T.B.), EU Marie Curie Career Integration Grant No. 303507, and German Research Foundation (DFG) Collaborative Research Centre (SFB) 1310 (to T.B.). A.E.C was supported by a Georg Forster fellowship from the Alexander von Humboldt Foundation.

## Author contributions

Conceptualisation: M.L. and T.B.; Investigation: M.L. and A.E.C.; Writing original draft: M.L.; Writing - review and editing: M.L., A.E.C., T.B.; Supervision: T.B.; Funding acquisition: T.B.

## Supplementary figures

**Figure S1:**
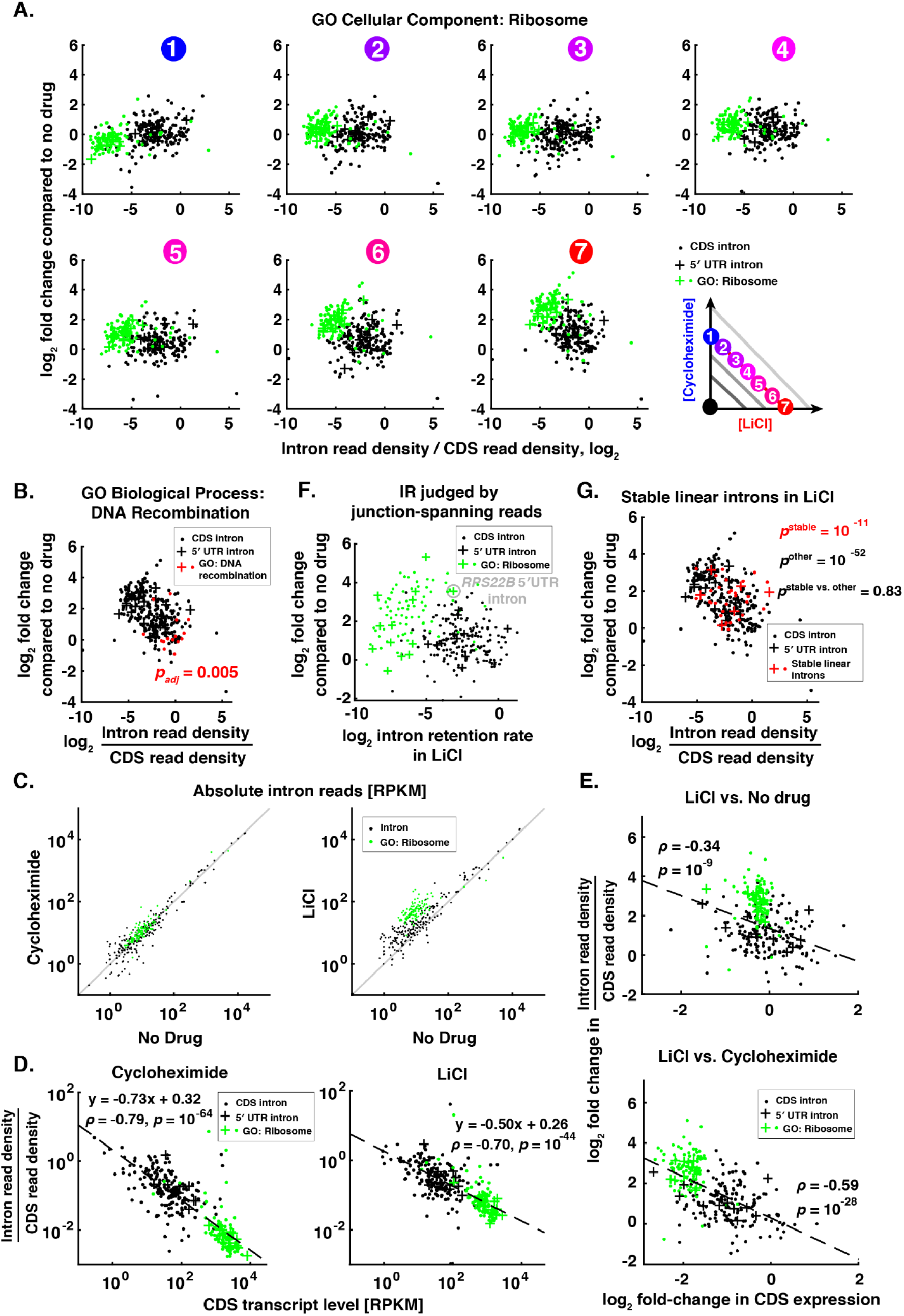
Introns in ribosomal proteins facilitate ribosomal protein response to LiCl stress.. **A.** Intron retention rates (as in Fig. 1C) along the 50% growth isobole of cycloheximide-LiCl drug combination. Introns contained in the genes belonging to the most significantly enriched GO Cellular Component with no offspring term as determined for intron retention increase in LiCl (‘Ribosome’) are highlighted in green. For quantification of intron retention rates from RNA sequencing data and gene enrichment analysis refer to *Methods*. **B.** Intron retention rate in LiCl (as in panel 7 in A). The introns from genes belonging to the GO Biological Process term most significantly enriched in the lower end of the relative increase in intron retention (‘DNA recombination’) is displayed in red, with the hypergeometric test *p*-value adjusted for multiple hypothesis testing. **C.** Intron read density plots for all introns, comparing different conditions, show that intron reads in LiCl are elevated with respect to the entire transcriptome, not just with respect to their parent transcript. Introns in ribosomal genes are highlighted in green. Grey lines are visual guides for no change. RPKM – reads per kilobase of transcript per million mapped reads. **D.** Intron retention in cycloheximide (left) or LiCl (middle) compared to no drug for individual introns negatively correlates with the expression level of the corresponding RNA as gauged by the RNA-seq read counts in coding region (CDS). Linear model was fitted over all introns. **E.** Change in intron retention in LiCl negatively correlates with change in the expression of the corresponding CDS, both when compared to no drug (top) and to cycloheximide (bottom). Linear model was fitted over all introns. **F.** Analysis of intron retention using only the sequencing reads overlapping the exon-intron junction as the proxy for intron retention rate (*Methods*). The good agreement with the analysis being performed using all reads in the introns (panel A) supports that introns are being retained within the transcripts rather than just being spliced and non-degraded. **G.** Stable linear introns previously implicated in starvation response are increasingly retained in LiCl just like the rest of the introns. *p*-values are reported for one-sided *t*-testing of whether the two intron groups (stable, other) are unchanged in LiCl; two-sided *t*-test of whether the two groups behave the same in LiCl shows no significant difference. All data in this figure are based on sequencing ribo-depleted total RNA.

**Figure S2:**
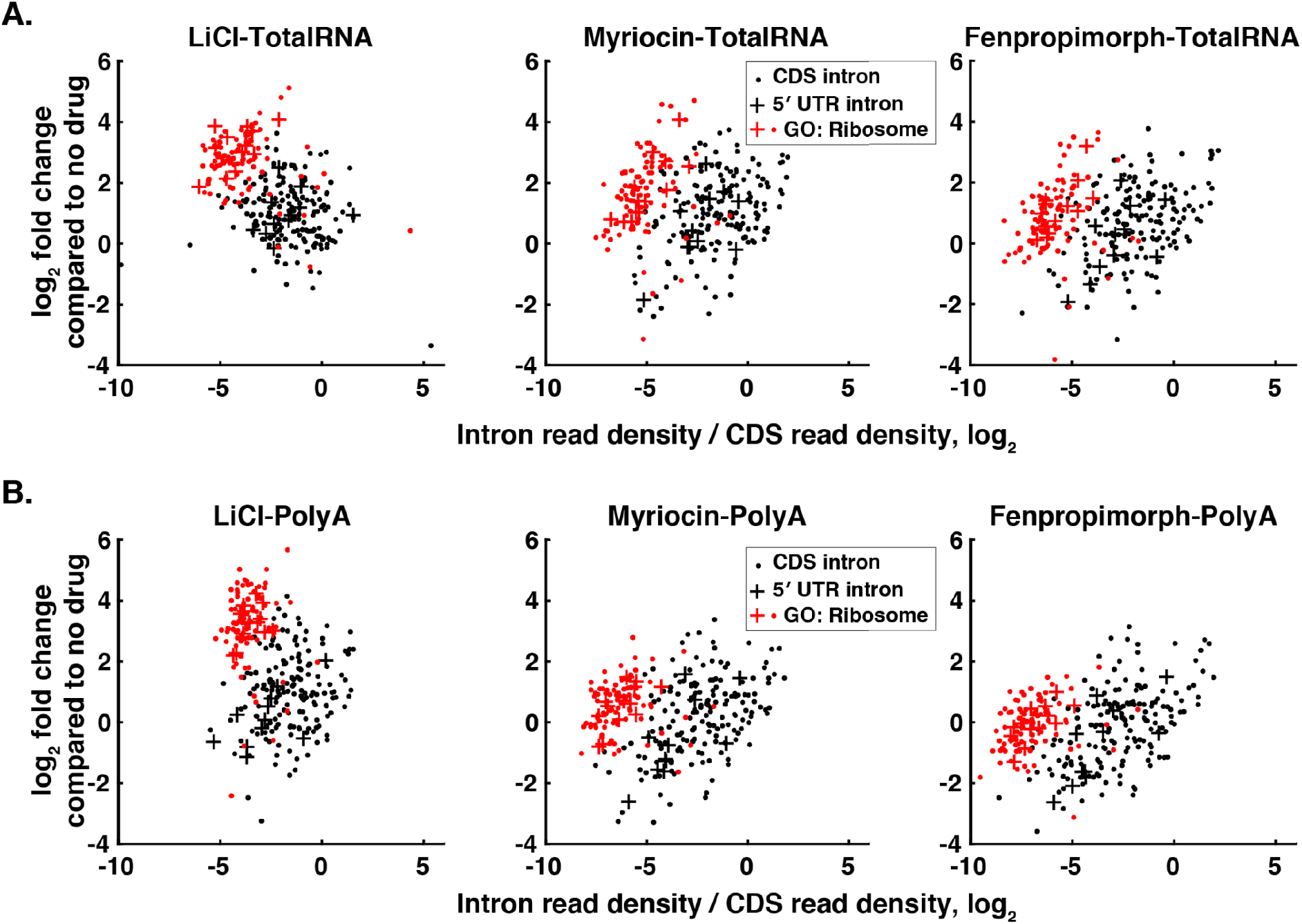
Sequencing of poly-A RNA fraction confirms that LiCl induces intron retention. **A.** Intron retention rate (intron read density / CDS read density) from total RNA sequencing, as in Fig. 1C, for cultures grown in the presence of the drugs shown at their IC_50_. **B.** As (A), but for polyA-sequencing assay.

**Figure S3:**
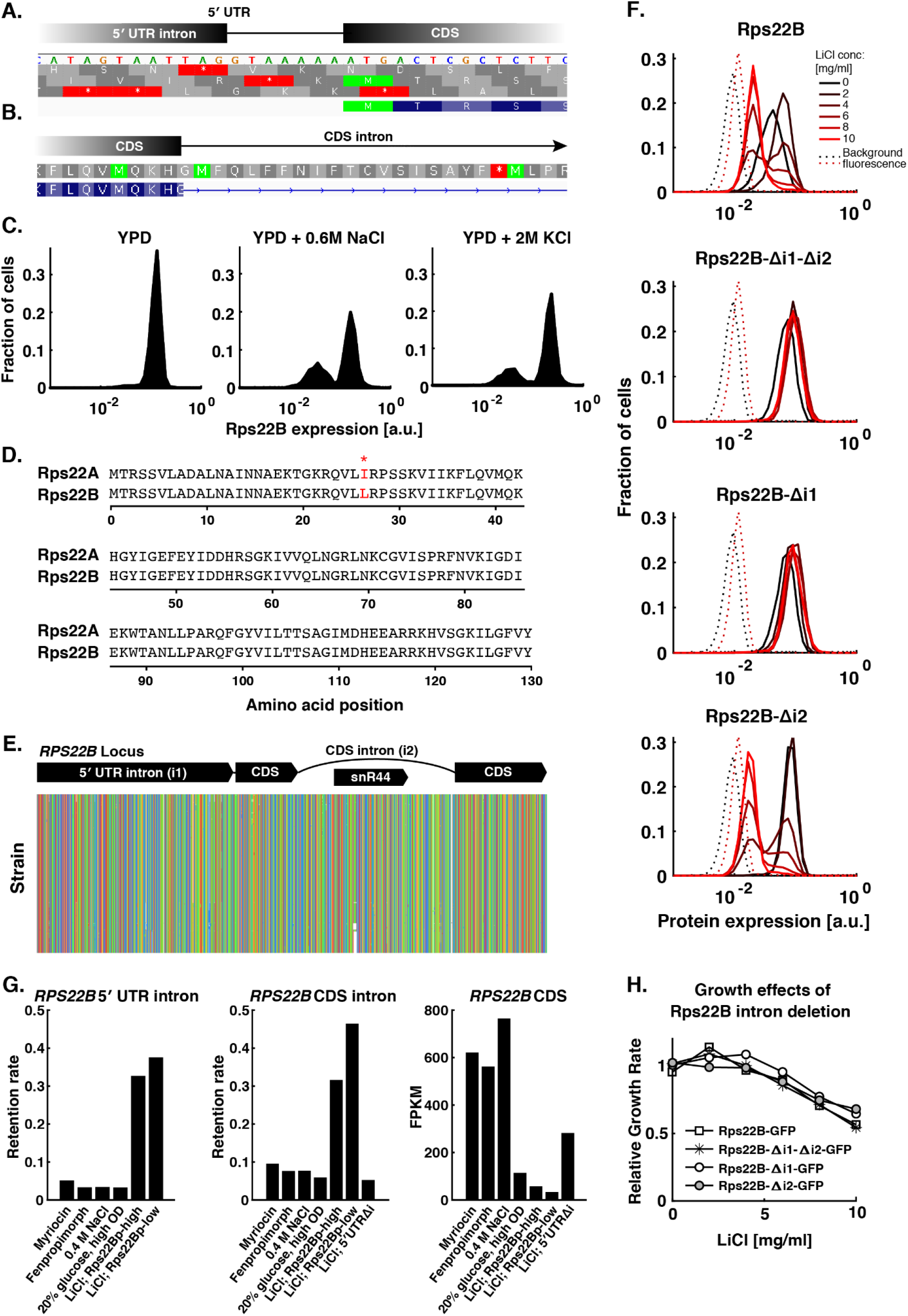
Osmotic stress induces bimodal expression in Rps22B. **A.** DNA sequence and translation reading frames around the 3’ splice site of the 5’ UTR *RPS22B* intron. Stop codons (red) are present in the vicinity in all three reading frames. **B.** Rps22B amino acid sequence and the corresponding translation reading frame at the 5’ splice site of *RPS22B* CDS intron, showing the presence of early stop codon in the intron. **C.** Histograms of Rps22B-GFP expression in NaCl and KCl, respectively, measured by flow cytometry during the exponential growth phase after 16 h of incubation. **D.** Protein blast for Rps22A and Rps22B. Red asterisk denotes the single difference between the amino acid sequences. RefSeq entries NP_013471.1 and NP_012345.1 are shown for Rps22A and Rps22B proteins, respectively. **E.** Multiple sequence alignment of previously published 1011 natural *Saccharomyces cerevisiae* isolates (*Methods*) reveals a remarkable conservation of the Rps22B 5’ UTR intron during within-species evolution. Rows represent individual isolates, colored columns represent individual bases in the Rps22B genomic locus (green=adenine, red=guanine, orange=cytosine, blue=thymine). **F.** Histograms of flow cytometry measurements of strains with GFP-tagged Rps22B, with seamless deletion of either or both introns in the presence of LiCl at different concentrations (*cf*. Fig. 2B). Δi1 denotes deletion of the 5’ UTR intron, Δi2 deletion of the CDS intron. **G.** Intron retention (intron read density / CDS read density) and transcript level (FPKM) for *RPS22B* measured by RNA sequencing of the polyA fraction in various conditions. Myricoin, fenpropimorph and LiCl were used at their respective IC_50_. For details on sorting of the Rps22B high and low subpopulations, see **Fig. 3G** and *Methods*. FPKM – fragments per kilobase per million reads, i.e. read density normalised to the number of reads in the sequencing run. **H.** Changes in growth rate due to intron deletion in Rps22B-GFP reporter strain cannot account for the observed differences in Rps22B expression pattern. Growth rates for the different strains are shown relative to the mean of their growth rates in the absence of drug.

**Figure S4:**
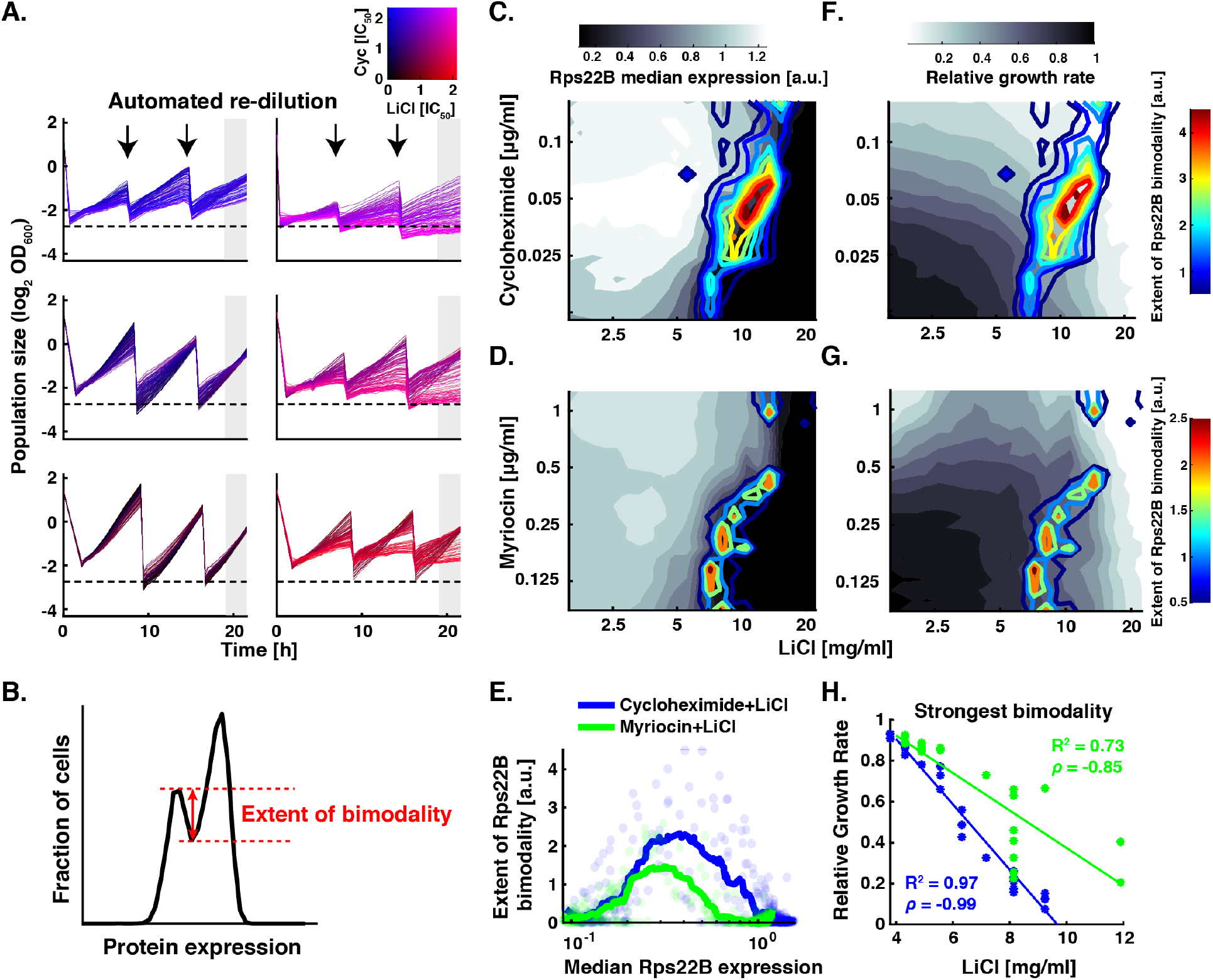
Rps22B protein expression is consistent with a bistable regulatory loop. **A.** Growth curves from the automated re-inoculation setup of cultures growing in a detailed 2-drug gradient of LiCl and cycloheximide, spread over 576 wells of six microtitre plates. Each panel shows growth curves for cultures on one 96-well plate. Drops in OD_600_ are due to automated re-dilution of growing cultures to an arbitrary target OD (dashed line). Shaded area denotes measurements that were used to determine the exponential growth rates shown in (F) and (G). **B.** Schematic showing the ad hoc definition of the quantity used to analyse the Rps22B bimodal expression pattern. The extent of bimodality is defined here as the y-axis distance between the trough and the lower of the two peaks on the histogram of single-cell protein expression, with bin size kept consistent throughout the analysis (*Methods*). **C. and D.** The extent of Rps22B bimodality (coloured lines), is overlaid on Rps22B median expression (greyscale) in a 2-drug gradient of LiCl and cycloheximide (C) or myriocin (D). **E.** Rps22B protein bimodality as a function of median Rps22B protein expression for all wells in the 2-drug gradient (dots) is shown along with a running average with a window of 30 data points (lines). Rps22B bimodality peaks at a certain level of median Rps22B expression, rather than at a certain growth rate or LiCl concentration. **F. and G.** The extent of Rps22B bimodality (coloured lines) is overlaid on growth rate (greyscale) in a 2D gradient of LiCl and cycloheximide (F) or myriocin (G). **H.** For each concentration of cycloheximide (blue) or myriocin (green), the LiCl concentration that elicited maximum Rps22B protein bimodality is plotted (x-axis), with the corresponding growth rate of the culture (y-axis). The strength of LiCl stress needed to bring the level of Rps22B to the putative unstable fixed point is inversely dependent on the growth rate, suggesting that the Rps22B switch is quantitatively attuned to specific growth conditions. *ρ* – Pearson correlation coefficient. See **Fig. S5** for Rps22B single-cell gene expression data for individual wells.

**Figure S5:**
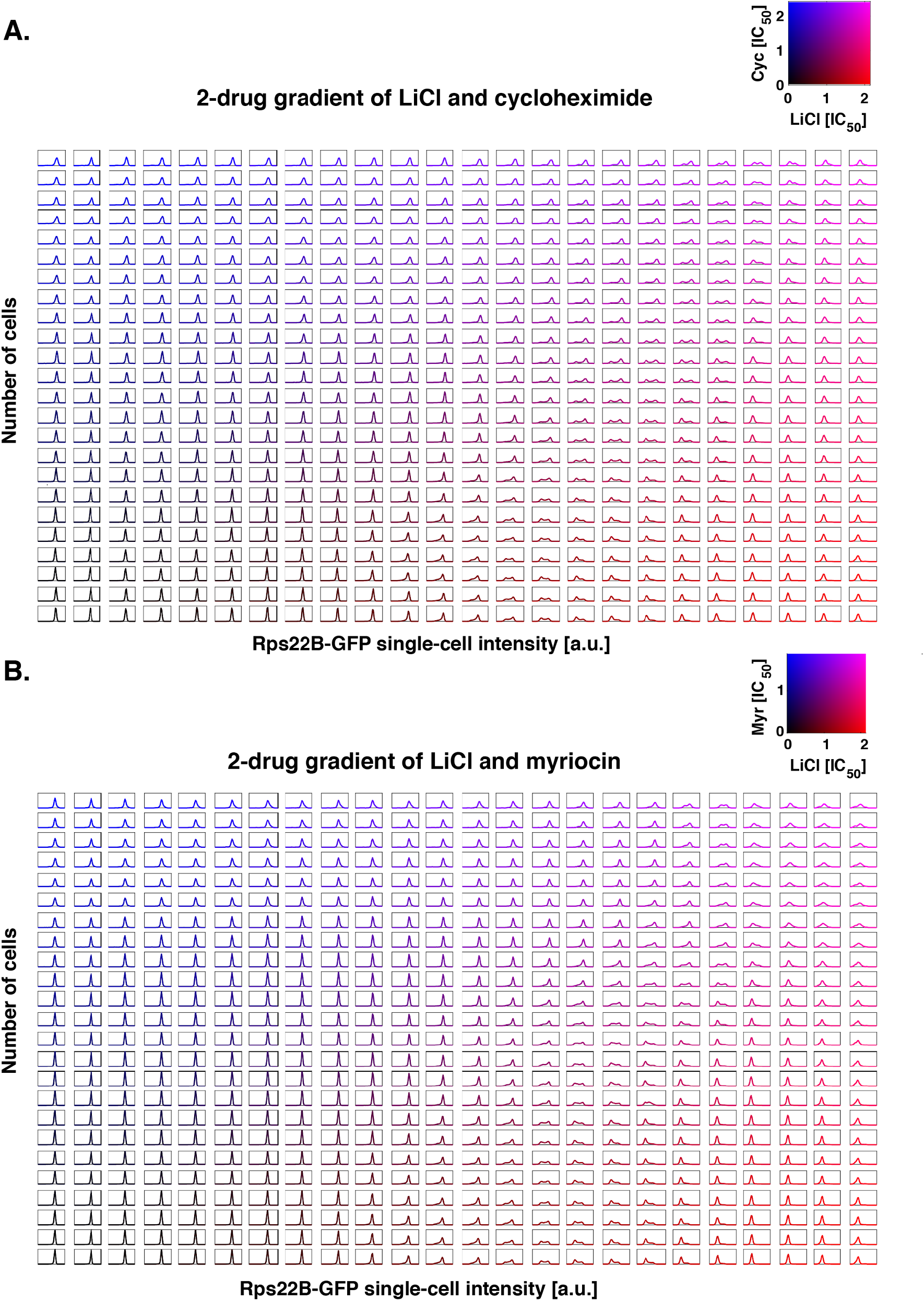
Rps22B single-cell protein expression in a detailed two-drug gradient. **A.** Histograms of flow cytometry measurements of strains with GFP-tagged Rps22B for 576 cultures with graded concentrations of LiCl and cycloheximide, incubated by automated reinoculation protocol (*Methods*). **B.** As (A), but for LiCl and myriocin.

**Figure S6:**
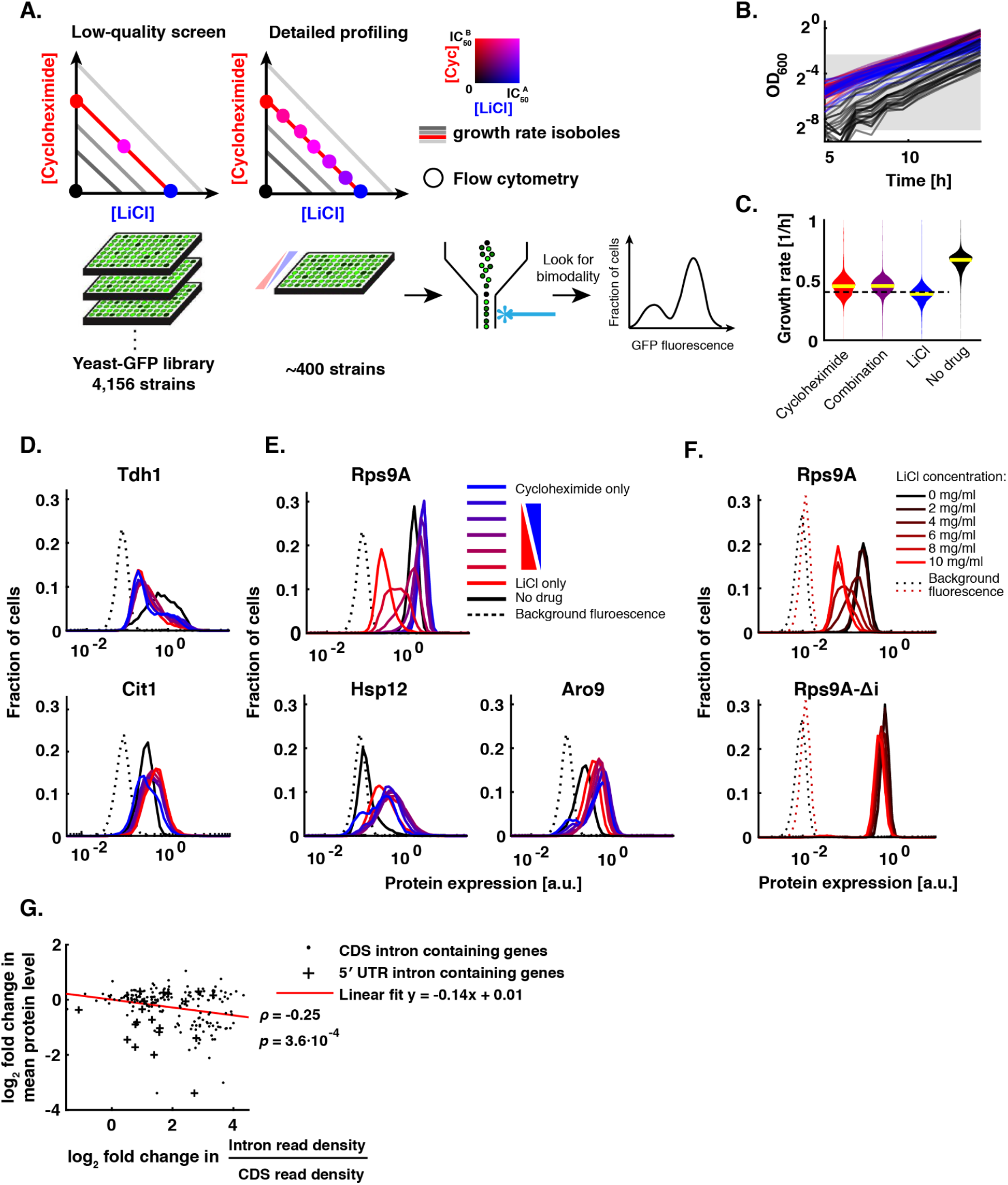
Single-cell isogrowth profiling uncovers drug-induced cellular states. **A.** Schematic showing the experimental strategy for single-cell isogrowth profiling. Using flow cytometry, the entire yeast protein-GFP library was screened in four conditions: no drug, each of the two drugs alone at 40% inhibition, and an equipotent combination of drugs at the same total inhibition. For about 10% of genes which showed an unusual expression pattern, the corresponding strains were restreaked and subjected to a more detailed screen in 8 conditions. **B**. Example growth curves of one 96-well plate with 24 strains each in 4 conditions. Note that the inoculum of the no-drug condition (black) was decreased in order to achieve comparable final cell density at the time of harvesting. The shaded area shows the OD range used for fitting of growth rates. **C.** Violin plot showing the distribution of growth rates for the initial screen over the entire library. The dashed line indicates 40% growth inhibition. **D.-E.** Single-cell GFP intensity histograms of protein-GFP fusion strains. Proteins with bimodal expression induced by growth inhibition by both LiCl and cycloheximide (D) or specifically by either drug (E) are shown. **F.** Intron deletion in Rps9A abolishes the LiCl-induced bimodality. **G.** Change in intron retention in LiCl compared to equal inhibition by cycloheximide as judged by RNA-sequencing is negatively correlated with the change in mean protein level as determined by flow cytometry. *ρ* – Pearson correlation coefficient; *p* value was determined by permutation test.

**Figure S7:**
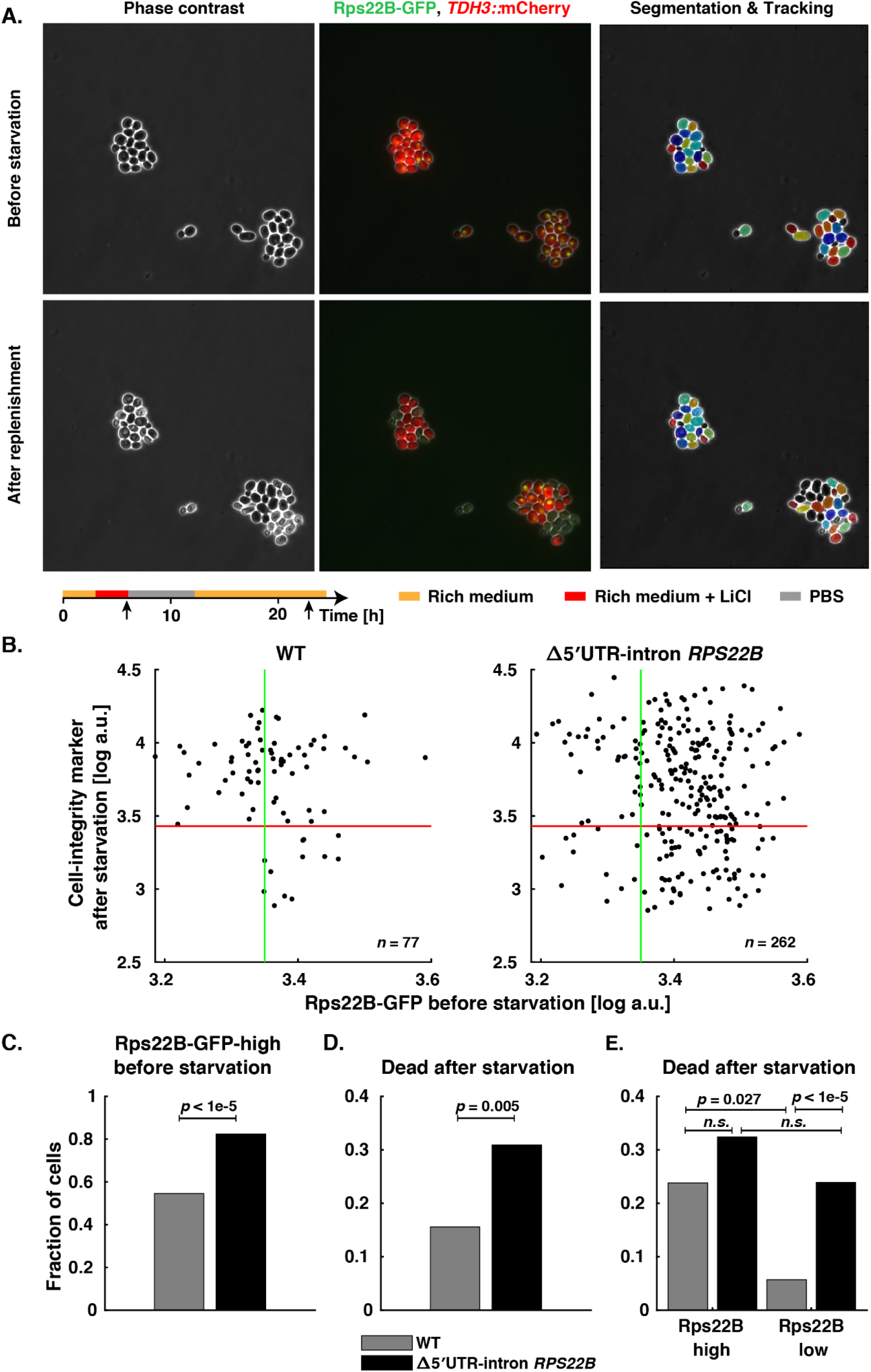
5’ UTR intron in *RPS22B* gene is necessary for induction of phenotypic heterogeneity by LiCl. **A.** Example micrographs of 5’ UTR intron-deleted Rps22B-GFP strain just before 6 h starvation in PBS (top row) and 10 h after rich medium replenishment (bottom row). Experimental setup is shown in the schematic below the micrographs; vertical arrows indicate timepoints at which the micrographs shown were taken. **B.** Quantification of mCherry (cell integrity marker) and GFP signal (*Methods*) in the micrographs from the timepoints in (A). **C**. 5’ UTR intron deletion in *RPS22B* increases the fraction of GFP-positive cells as defined by the green lines in B. **D.** Same as Fig. 3K: Fraction of dead cells after starvation for WT (grey) and 5’ UTR intron-deletion mutant (black). 5’ UTR intron deletion in *RPS22B* increases the fraction of cells that do not survive the starvation stress, determined based on mCherry expression. The mCherry threshold for dead cells shown in (B) was chosen to be lower than mCherry expression of any cell before the starvation as well as lower than that of any cell after starvation that resumed growth. **E.** As (D), but showing Rps22B-high and -low subpopulations separately. The 5’ UTR intron deletion in *RPS22B* largely abolishes the coupling of the phenotypic heterogeneity in cell death to Rps22B protein expression heterogeneity. The *p*-values were determined using a permutation test; *n.s.* – not significant, *p* > 0.05.

**Figure S8:**
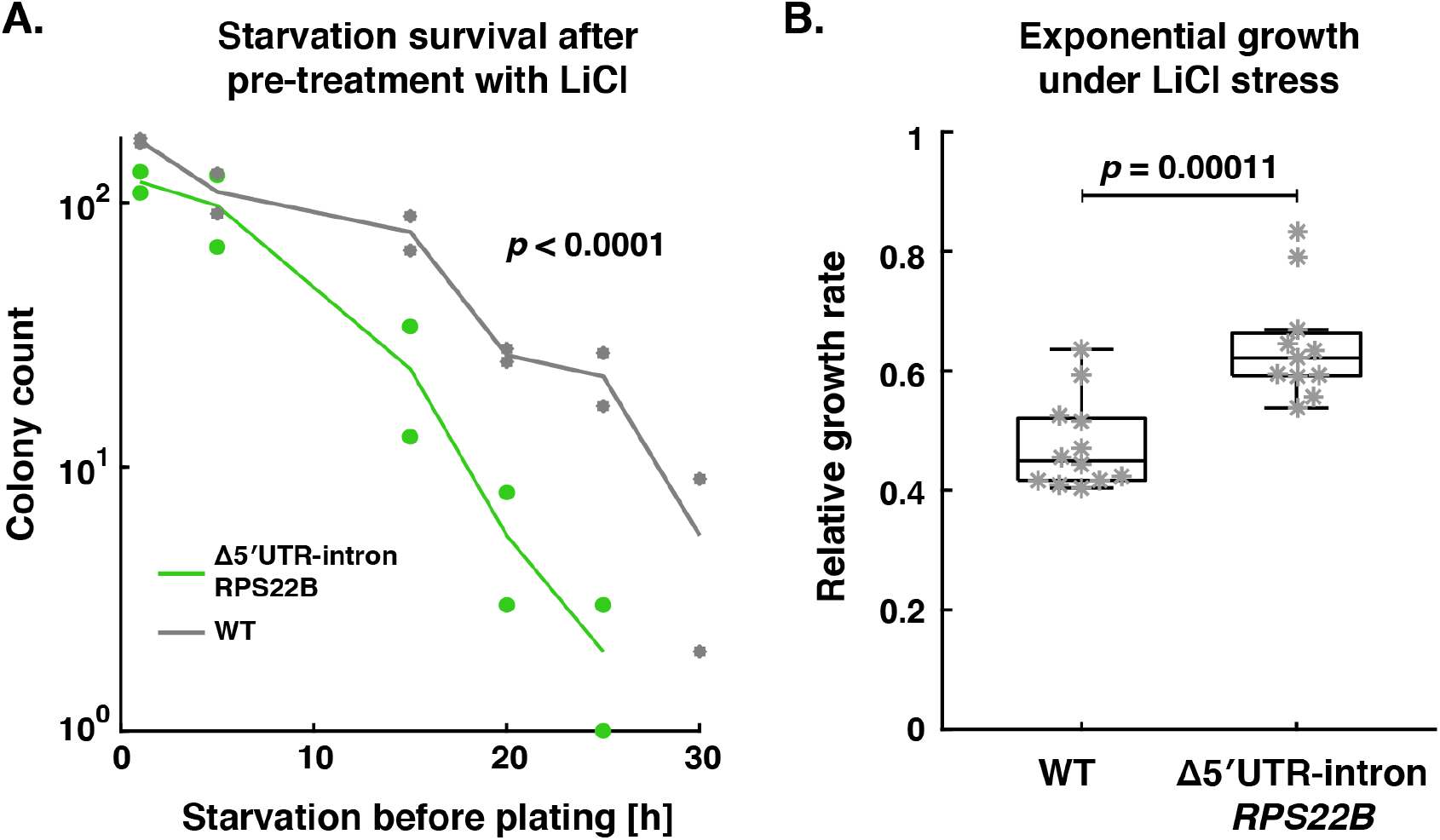
Deletion of 5’ UTR intron in *RPS22B* confers phenotypic differences under starvation and osmotic stress. **A.** Survival curves of the intron deletion mutant (green) and WT (grey), pre-cultured in YPD containing 9 mg/ml LiCl, when starved in PBS for varying duration before being plated on rich medium; pre-culturing the WT strain in LiCl at 9mg/ml results in a uniform population with low Rps22B expression (Fig. 2A), while deletion of the 5’ UTR intron results in a uniform population with high Rps22B expression (Fig. 2B). Significance was determined as in Fig. 3H. **B.** Exponential growth rates relative to uninhibited WT, of the 5’ UTR intron-deletion mutant and WT strain, in YPD containing 9 mg/ml LiCl. Two-sided *t*-test was used to determine significance.

**Figure S9:**
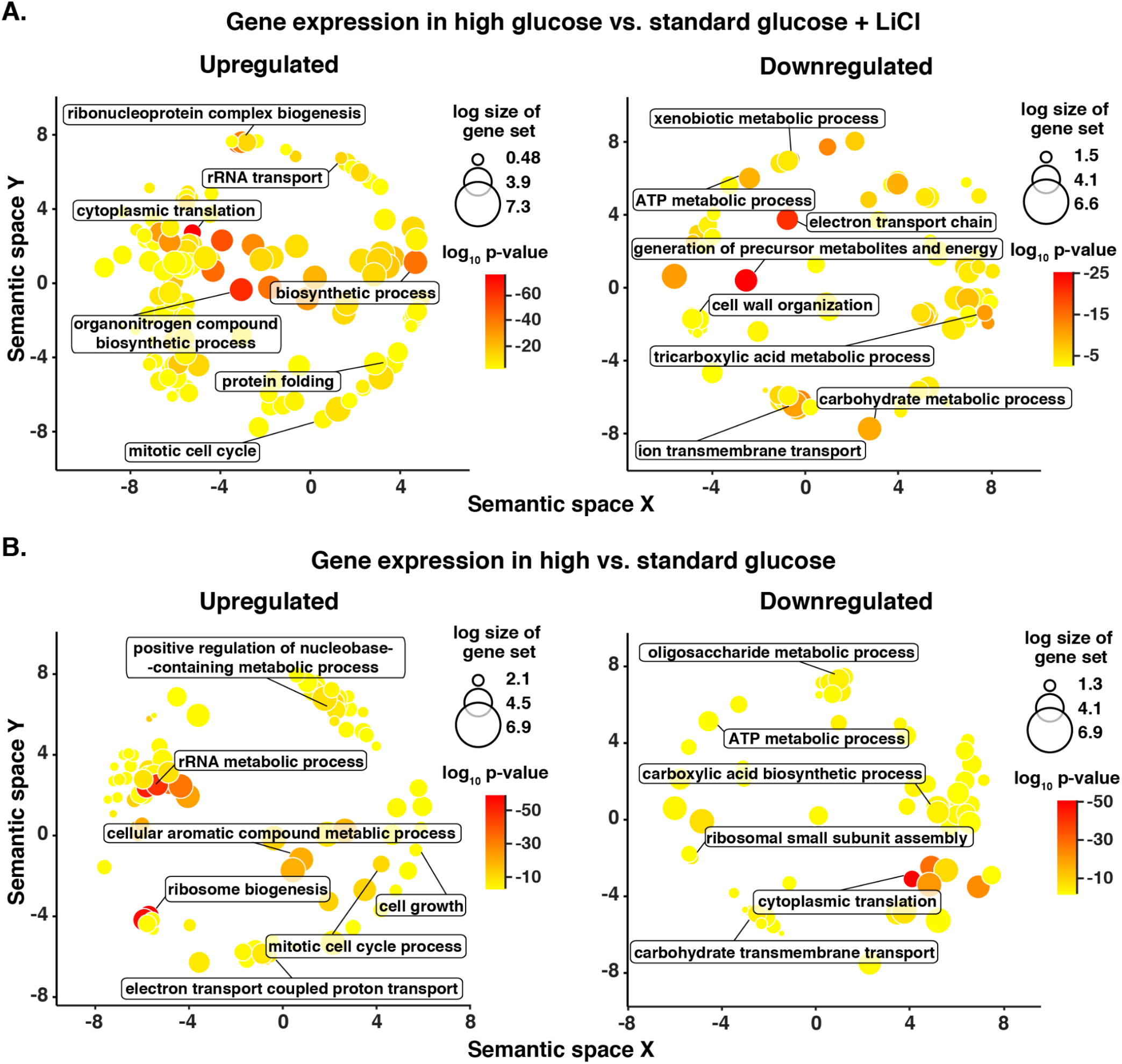
Functional enrichment analysis of gene expression changes in hyperosmotic glucose. Revigo visualisations (*Methods*) of GO:Biological Process enrichment analysis of total RNA sequencing reads from a yeast culture grown in 20% glucose to late exponential phase compared to a culture grown to mid-exponential phase in standard YPD containing IC_50_ LiCl (A) or no drug (B). GO terms significantly upregulated (left) or downregulated (right) in high glucose are shown, along with GO term names for selected representatives of individual clusters.

## Supplementary movies

https://drive.google.com/file/d/1Ezd7Awe3qpvzDla-fezjaim-ievUtGFQ/view?usp=sharing

**Supplementary Movie 1: Long starvation favours Rps22B-low cells in that they lyse less frequently than Rps22B-high cells.**

Time lapse microscopy video of Rps22B-GFP yeast strain grown in a microfluidic chamber, flushed with medium as described in **Fig. 3A** left panel. Size bar = 10 μm.

https://drive.google.com/file/d/1vksbIdPIOHgSXJGycNfbn34G2zWuYjom/view?usp=sharing

**Supplementary Movie 2: Short starvation favours Rps22B-high expressing cells in that they start growing faster after the replenishment of nutrients.**

Time lapse microscopy video of Rps22B-GFP yeast strain grown in a microfluidic chamber, flushed with medium as described in **Fig. 3A** right panel. Size bar = 10 μm.

## Supplementary tables

https://drive.google.com/file/d/1qe-cg6THpaOq7Fj4swhO2ZGfonxBniXn/view?usp=sharing

**Table S1: Flow cytometry summary data for strains from the yeast protein-GFP library profiled in four conditions.**

For each profiled strain and condition, the summary statistics for FITC-H channel normalized cell-wise to FSC-H channel is listed, including arithmetic mean, standard deviation, and relative counts in 100 histogram bins spaced logarithmically from 10^-3^ to 10^5^. The strains that were profiled in the more detailed 8-point measurement (**Table S2**), are not listed.

https://drive.google.com/file/d/1l2MQfiM3ortSvDcWhcskqxDrz5NWXQKx/view?usp=sharing

**Table S2: Flow cytometry summary data for strains from the yeast protein-GFP library profiled in eight conditions.** For each profiled strain and condition, the summary statistics for FITC-H channel normalized cell-wise to FSC-H channel is listed, including arithmetic mean, standard deviation, and relative counts in 100 histogram bins spaced logarithmically from 10^-3^ to 10^5^. The measurement of parent BY4741 (non-GFP) strain is listed as well for background fluorescence reference.

## Notes

### Competing Interest Statement

The authors have declared no competing interest.

